# Identification of gene products involved in plant colonization by *Pantoea* sp. YR343 using a diguanylate cyclase expressed in the presence of plants

**DOI:** 10.1101/2021.03.03.433726

**Authors:** Amber N. Bible, Mang Chang, Jennifer L. Morrell-Falvey

## Abstract

Microbial colonization of plant roots is a highly complex process that requires the coordination and regulation of many gene networks, yet the functions of many of these gene products remain poorly understood. *Pantoea* sp. YR343, a gamma-proteobacterium isolated from the rhizosphere of *Populus deltoides*, forms robust biofilms along the root surfaces of *Populus* and possesses plant growth-promoting characteristics. The mechanisms governing biofilm formation along plant roots by bacteria, including *Pantoea* sp. YR343, are not fully understood and many genes involved in this process have yet to be discovered. In this work, we identified three diguanylate cyclases in the plant-associated microbe *Pantoea* sp. YR343 that are expressed in the presence of plant roots, One of these diguanylate cyclases, DGC2884 localizes to discrete sites in the cells and its overexpression results in reduced motility and increased EPS production and biofilm formation. We then performed a genetic screen by expressing this diguanylate cyclase from an inducible promoter in order to identify candidate downstream effectors of c-di-GMP signaling which may be involved in root colonization by *Pantoea* sp. YR343. Further, we demonstrate the importance of other domains in DGC2884 to its activity, which in combination with the genes identified by transposon mutagenesis, may yield insights into activity and regulation of homologous enzymes in medically and agriculturally relevant microbes.

## Introduction

The ability of plant growth promoting bacteria to exert beneficial effects on plant hosts is mediated through chemical and physical associations with plant tissues. Associating with the plant and surviving within this environment likely requires the coordination of multiple signaling pathways. For example, chemotaxis signaling pathways are involved in the detection of chemicals found in plant root exudates (1–4). Moreover, plants are capable of tailoring the composition of root exudate to promote associations with specific microbes (5). Many bacterial species also use quorum-sensing as a mechanism in the rhizosphere for influencing changes in gene expression that can lead to root colonization and biofilm formation (6–9). Indeed, genome analyses showed that acyl-homoserine lactone (AHL)-based signaling systems are prevalent in the microbiome of *Populus deltoides* (10). Additionally, plant colonization involves the second messenger signaling molecule, cyclic diguanylate monophosphate (c-di-GMP), which is known to affect motility, virulence, exopolysaccharide (EPS) production, and biofilm formation in many bacterial species (11–15).

The levels of c-di-GMP within cells are regulated by two different enzymes: diguanylate cyclases, which catalyze the production of c-di-GMP from two molecules of guanosine triphosphate (GTP), and phosphodiesterases, which degrade c-di-GMP to guanosine monophosphate (GMP) (11–13, 16). Most bacterial genomes encode many diguanylate cyclases and phosphodiesterases, suggesting that control of c-di-GMP levels is highly regulated. Production of c-di-GMP by diguanylate cyclases has been shown to modulate a wide variety of cellular behaviors through different types of effector proteins. For instance, the flagellar brake protein, YcgR, from *E. coli* and *Salmonella enterica* serovar Typhimurium, can bind c-di-GMP through a PilZ domain in order to modulate motility (17, 18), while the transcriptional regulator, VpsT, regulates biofilm formation in *V. cholera* in response to c-di-GMP levels (19). Thus far, c-di-GMP has been shown to bind to proteins containing PilZ or GIL domains (20–22), as well as riboswitch proteins (23). Furthermore, c-di-GMP has also been shown to bind to the RxxD I-sites of many diguanylate cyclases and those with degenerate GGDEF domains often serve as effector proteins (24).

Prior studies have shown that diguanylate cyclases can be regulated at either the level of enzymatic activity or the level of expression, based on conditions within the surrounding environment (11). Here, we wanted to identify diguanylate cyclases that were regulated at the expression level in response to the presence of a plant in the rhizosphere using *Pantoea* sp. YR343. *Pantoea* sp. YR343 was isolated from the rhizosphere of *Populus deltoides* and has been shown to associate with a variety of plant hosts, including *Populus deltoides*, *Populus trichocarpa, Triticum aestivum*, and *Arabidopsis thaliana* (25, 26). *Pantoea* sp. YR343 is a gamma-proteobacterium from the *Enterobacteriaceae* family which produces indole-3-acetic acid (IAA) (27, 28) and solubilizes phosphate, both of which have been shown to contribute to plant growth (6). Furthermore, *Pantoea* sp. YR343 has been shown to form biofilms along the surface of plant roots (25). Because c-di-GMP plays an important role in biofilm formation, we hypothesized that there may be diguanylate cyclases that are specifically expressed in response to growth in the presence of a plant root in the rhizosphere. To this end, we identified three diguanylate cyclases that are expressed during colonization of plant roots. Overexpression of one of these diguanylate cyclases (encoded by PMI39_02884 and hereby referred to as DGC2884) significantly impacted EPS production, motility, and biofilm formation. This overexpression strain was utilized for a genetic screen to identify candidate genes that affect the ability of *Pantoea* sp. YR343 to regulate EPS production in the presence of high levels of c-di-GMP, which we hypothesized would have defects in biofilm formation and root colonization. Transposon mutants affecting several of these genes were further characterized for their ability to colonize plant roots.

## Results

### Diguanylate cyclase promoter analysis

The genome of *Pantoea* sp. YR343 contains 13 genes predicted to encode proteins with diguanylate cyclase domains, 8 genes predicted to encode phosphodiesterase domains, 8 genes predicted to encode proteins possessing both diguanylate cyclase and phosphodiesterase domains, and 3 genes predicted to encode proteins with PilZ-domains (https://img.jgi.doe.gov). Table 1 lists each of the diguanylate cyclases found in *Pantoea* sp. YR343, along with its domain organization based on Pfam analyses (29). Notably, 13 of the 21 predicted diguanylate cyclases lack a canonical RxxD I-site (Table 1). We hypothesized that of the 21 predicted diguanylate cyclases, there were likely some enzymes that were expressed in response to the presence of a plant in the rhizosphere. In order to identify candidate enzymes, we began by generating promoter-reporter constructs for each of the 21 genes encoding diguanylate cyclase domains by fusing each promoter to the gene encoding green fluorescent protein (GFP) using a pPROBE-NT vector (30). Putative gene promoters for each enzyme were predicted using BPROM (31). We were able to successfully produce these promoter fusion constructs for 20 of the 21 diguanylate cyclases (Table 1). After transforming these constructs into wild type *Pantoea* sp. YR343, the reporter strains were grown under different conditions to determine under which growth conditions the promoters were active, compared to a control strain carrying an empty pPROBE-NT vector. We measured the average fluorescence intensity of cells grown under various growth conditions, and then normalized fluorescence intensity values against the empty vector control strain set to a value of 1.00. When cells were grown in M9 minimal media with 0.4% glucose, we found that twelve diguanylate cyclase reporters showed an average fluorescence intensity below 2.00 (weak or no expression), making them suitable candidates for further study in terms of expression in biofilms, pellicles, and during root colonization (Table 1). To test for expression during biofilm formation, the cells were grown statically in M9 minimal medium with 0.4% glucose for 72 hours in 12-well dishes containing a vinyl coverslip as described in Materials and Methods. These data show that eleven diguanylate cyclases showed increased expression under these conditions, with DGC2884 and DGC2242 showing the highest levels (Table 1 and Fig 1). Interestingly, we found that each of the strains showed an increase in expression during biofilm formation based on GFP fluorescence, but images showed that GFP levels driven from the DGC2884 promoter were not uniform within the biofilm (Fig 1). Instead, we found that GFP was highly expressed in specific patches throughout the biofilm, but expressed at low or undetectable levels in other regions. This expression pattern was also observed in some of the other promoter constructs and is reflected, in part, by the higher S.E.M. values shown in Table 1. We also tested for expression during pellicle formation and found that most strains only exhibited a modest increase in expression (Table 1).

**Figure 1.**
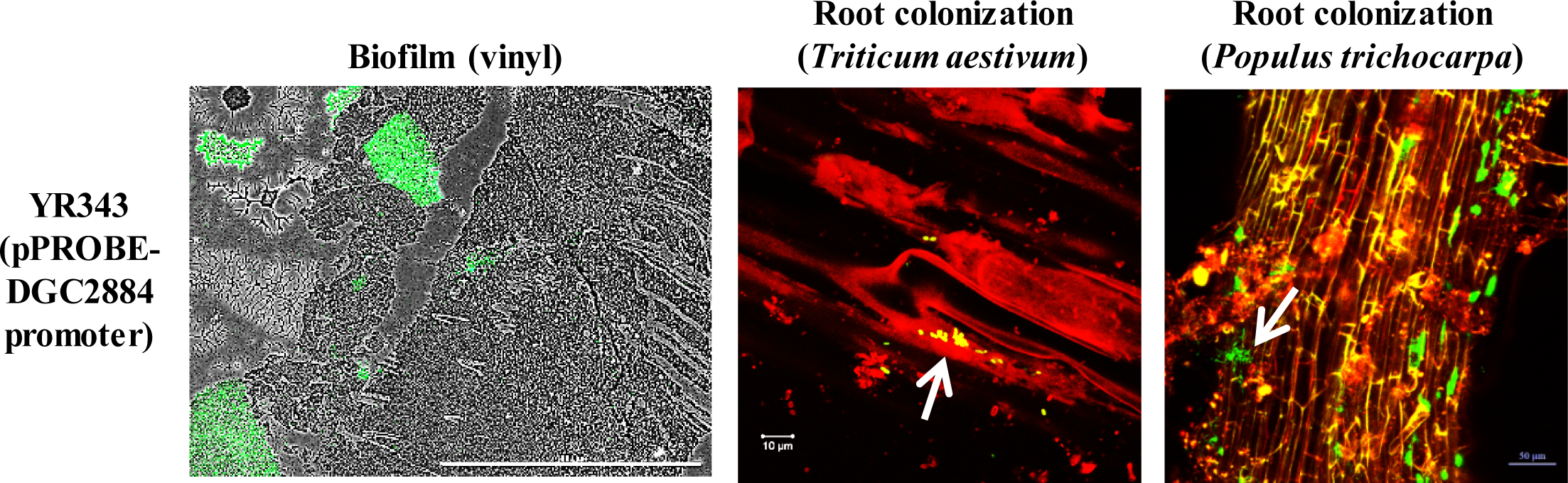
Promoter-GFP reporter assays for DGC2884 expression under different growth conditions: biofilms on vinyl coverslips, *T. aestivum* root colonization and *P. trichocarpa* root colonization. Scale bars represent 1 mm in biofilm image, while scale bars in root colonization images are labeled accordingly. Arrows indicate bacterial colonization along the surface of plant roots. Data is representative of a minimum of three independent experiments.

**TABLE 1.** Promoter activity under various growth conditions

| DGC<sup>1</sup> | Gene Locus Tag | Domain architecture<sup>2</sup> | liquid culture | biofilm | pellicle | root colonization |
| --- | --- | --- | --- | --- | --- | --- |
| pPROBE | control |  | 1.00 | 1.00 | 1.00 | - |
| DGC2884 | PMI39_02884 | CHASE8-GGDEF | 1.16 ± 0.18 | 1841.20 ± 938.14 | 1.28 ± 0.17 | + |
| DGC3006 | PMI39_03006 | PAS3-GGDEF | 1.05 ± 0.11 | 105.44 ± 18.06 | 1.08 ± 0.18 | + |
| DGC3134 | PMI39_03134 | MASE5-GGEEF | 0.75 ± 0.08 | 11.76 ± 3.16 | 1.19 ± 0.44 | + |
| DGC0751 | PMI39_00751 | dCache_1-GGEEF (RxxD) | 0.95 ± 0.12 | 0.90 ± 0.43 | 1.03 ± 0.16 | - |
| DGC0995 | PMI39_00995 | GGEEF | 0.95 ± 0.02 | 5.09 ± 2.37 | 0.90 ± 0.09 | - |
| DGC1023 | PMI39_01023 | MHYT-MHYT-MHYT-GGDEF-EAL | 1.62 ± 0.03 | 217.99 ± 35.02 | 1.21 ± 0.18 | - |
| DGC1024 | PMI39_01024 | MHYT-MHYT-MHYT-GGDEF-EAL | 0.78 ± 0.02 | 67.83 ± 28.19 | 0.98 ± 0.05 | - |
| DGC2242 | PMI39_02242 | CHASE4-GGDEF-EAL (RxxD) | 0.87 ± 0.11 | 1318.75 ± 112.66 | 0.93 ± 0.10 | - |
| DGC3247 | PMI39_03247 | GGEEF (RxxD) | 0.94 ± 0.03 | 24.94 ± 8.30 | 0.95 ± 0.13 | - |
| DGC3482 | PMI39_03482 | GAF-GGDEF (RxxD) | 0.94 ± 0.02 | 38.13 ± 9.60 | 1.20 ± 0.14 | - |
| DGC3621 | PMI39_03621 | dCache_1-GGEEF (RxxD) | 1.27 ± 0.01 | 22.88 ± 14.83 | 1.42 ± 0.12 | - |
| DGC0366 | PMI39_00366 | GAPES4-FRSDF-ELL | 5.09 ± 0.69 | N. D.<sup>3</sup> | N. D. | N. D. |
| DGC1008 | PMI39_01008 | PAS9-PAS9-PAS9-PAS4-GGDEF-EAL (RxxD) | 4.35 ± 0.11 | N. D. | N. D. | N. D. |
| DGC1089 | PMI39_01089 | GGEEF | 4.31 ± 0.11 | N. D. | N. D. | N. D. |
| DGC1854 | PMI39_01854 | MASE1-PAS3-PAS3-PAS-GGDEF (RxxD) | 20.95 ± 0.76 | N. D. | N. D. | N. D. |
| DGC2196 | PMI39_02196 | CHASE7-GGEEF (RxxD) | 5.33 ± 0.03 | N. D. | N. D. | N. D. |
| DGC2334 | PMI39_02334 | GAPES4-GPSDF-EAL | NA<sup>4</sup> | N. D. | N. D. | N. D. |
| DGC2465 | PMI39_02465 | CHASE4-PAS-GGDAF-EAL | 2.00 ± 0.03 | N. D. | N. D. | N. D. |
| DGC2697 | PMI39_02697 | PAS9-GGDEF-EAL | 16.73 ± 0.27 | N. D. | N. D. | N. D. |
| DGC3217 | PMI39_03217 | GGEEF | 12.77 ± 0.81 | N. D. | N. D. | N. D. |
| DGC4070 | PMI39_04070 | MASE2-GGDEL | 7.76 ± 0.18 | N. D. | N. D. | N. D. |

Promoter assay performed using pPROBE vector cloned into *Pantoea* sp. YR343 and grown under different conditions. Values for average fluorescence (along with the standard deviation) are reported here. See Materials and Methods for details.
<sup>1</sup>Diguanylate cyclase gene for which the promoter was assayed for activity.
<sup>2</sup>Domain architecture and nomenclature is based on sequence analysis using the Pfam database. (RxxD) indicates that this enzyme possesses a RxxD site.
<sup>3</sup>N. D., not determined
<sup>4</sup>NA, not available
<sup>5</sup>Fluorescence was described in qualitative terms, where “+” indicates observed fluorescence and “-” indicates no observed fluorescence.

Next, we tested the activity of these 12 promoters during root colonization of *T. aestivum and P. trichocarpa*. Bacteria associated with roots were examined for the presence or absence of fluorescence, since quantification of expression levels was difficult due to plant autofluorescence (Table 1). After one week of growth post-inoculation, we found that DGC2884, DGC3006, and DGC3134 were expressed on *T. aestivum* and *P. trichocarpa* roots (Fig 1 and Table 1). We cannot exclude the possibility that the eight untested diguanylate cyclases may also be expressed during plant association since their high levels of background fluorescence during growth in liquid culture precluded testing them directly on plants. For the purpose of this study, however, we chose to focus on one of the three enzymes that were expressed during root colonization which was DGC2884.

### Characterization of DGC2884 and mutant variants in biofilm formation, Congo Red binding, and motility

The domain architecture of DGC2884 lacks a RxxD I-site, and the N-terminal CHASE8 domain of DGC2884 consists of a transmembrane domain and a HAMP (Histidine kinase, Adenylate cyclase, Methyl-accepting protein, and Phosphatase) domain (Table 1). In the absence of an I-site, the two glycine residues in the GGDEF domain are essential for binding of c-di-GMP (32). Interestingly, this domain architecture is not altogether unusual and has been studied in other enzymes. The best studied diguanylate cyclases with these domains are YfiN (or TpbB) from *Pseudomonas aeruginosa*, *Salmonella enterica*, and *Escherichia coli* (also called DgcN) (17, 33–35). These enzymes have been shown to primarily regulate motility via YcgR and production of exopolysaccharides (such as Pel and Psl in *P. aeruginosa*), in addition to roles in cell division (17, 33–35). Among these examples, DgcN from *E. coli* is the only example to lack a RxxD I-site, like DGC2884 from *Pantoea* sp. YR343. Interestingly, multiple sequence alignments using protein sequences in Clustal Omega show a 33% sequence identity to TpbB from *P. aeruginosa* and 37% sequence identity to DgcN from *E. coli* (Fig S1) (36). Furthermore, YfiN in *P. aeruginosa* and *E. coli* is found within the YfiBNR operon, which consists of the outer membrane lipoprotein YfiB, which stimulates YfiN, and the soluble periplasmic protein YfiR, which represses the activity of YfiN (33, 34). Interestingly, DGC2884 is located within an operon that resembles the YfiBNR operon, suggesting that DGC2884 may have a similar function to YfiN. In recent work, it has been suggested that in the absence of a RxxD I-site, the transmembrane and HAMP domains work to dimerize the diguanylate cyclase and allow it to bind to two c-di-GMP molecules (32). We therefore hypothesized that loss of the N-terminal transmembrane domain is critical to the function of DGC2884 in *Pantoea* sp. YR343.

To further characterize the diguanylate cyclase DGC2884 from *Pantoea* sp. YR343, we generated constructs with the full length DGC2884, an enzyme-dead DGC2884 AADEF mutant, and a DGC2884ΔTM mutant and overexpressed each of these in a wild type *Pantoea* sp. YR343 background. Construction and characterization of DGC2884 with the GGDEF motif mutated to AADEF, which has been shown to render diguanylate cyclases enzymatically inactive (32, 37–41), was used to further support that enzyme activity of this enzyme was responsible for any observed phenotypes. We next examined how expression of DGC2884 and its variants affected colony morphology, Congo Red binding, biofilm formation, and motility in comparison to a control strain carrying an empty vector (Fig 2). Growth curves were compared in both minimal and rich media (Fig S2). Notably, expression of wild type DGC2884, but not any of the variants, resulted in formation of small aggregates, likely skewing the measurements of optical density (data not shown). In prior work, we found that *Pantoea* sp. YR343 exhibits drier colony morphology on LB media and a more mucoid phenotype on R2A media (25); therefore, we first examined growth of these strains on each media type containing Congo Red, a dye specific to β-linked glucans and curli fibers (42). These results show that *Pantoea* sp. YR343 cells overexpressing DGC2884 (YR343 (pSRK (Km)-*DGC2884*)) resulted in red, wrinkly colony formation (Fig 2A). In contrast, overexpression of the AADEF variant (YR343 (pSRK (Km)-*DGC2884 AADEF*)) resulted in a colony phenotype more similar to the empty vector control, with the exception of wrinkles, suggesting that expression of DGC2884 in the absence of enzymatic activity may still retain some function (Fig 2A). We observed that Congo Red binding by strains expressing the DGC2884ΔTM variant was less than that of the DGC2884 expressing strain and we no longer observed wrinkly colony morphology, supporting the hypothesis that the TM domain of DGC2884 is critical to its function (Fig 2A).

**Figure 2.**
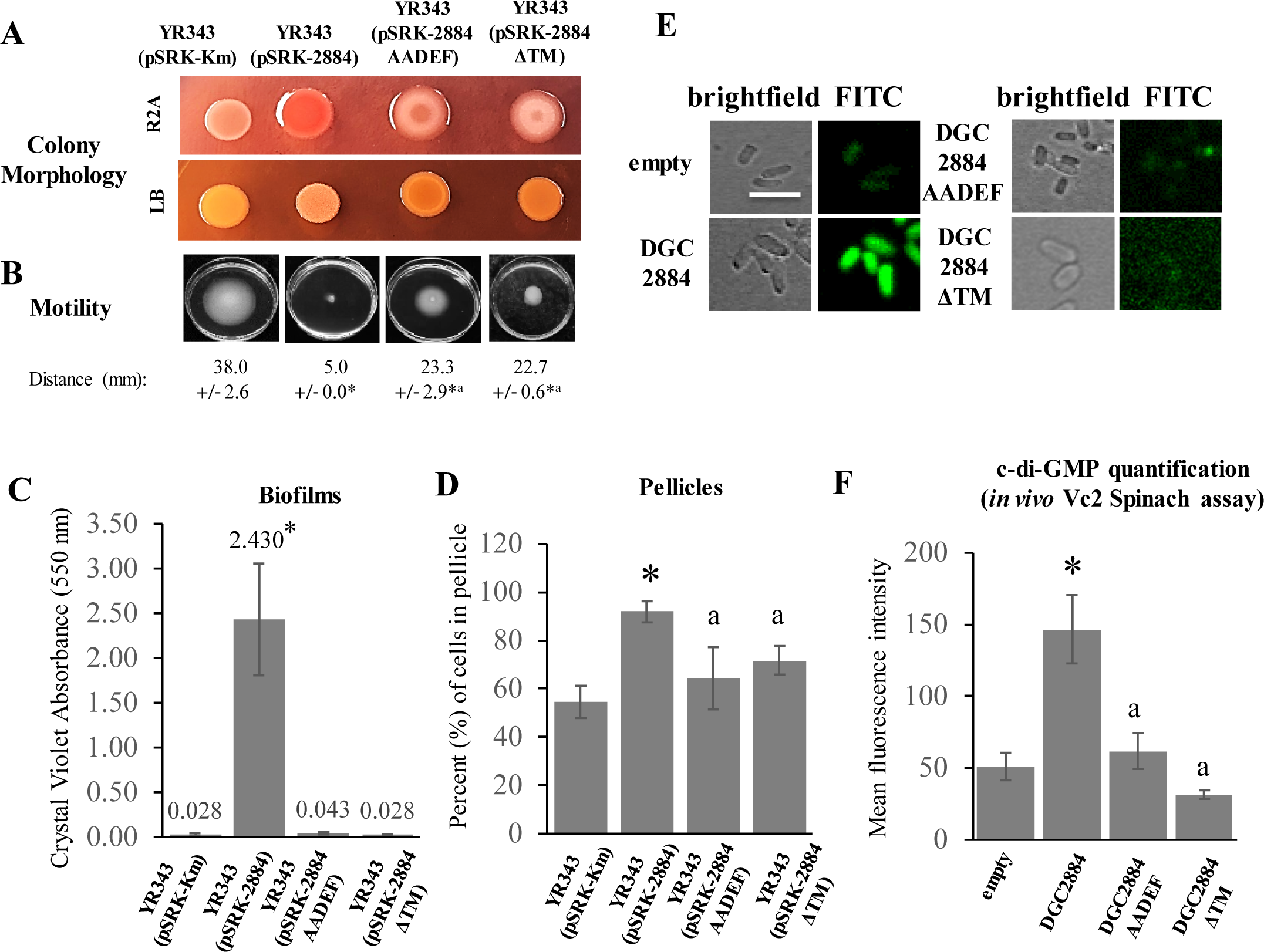
Characterization of individual diguanylate cyclases expressed in a wild type *Pantoea* sp. YR343 background. (A) Indicated strains were spotted on either LB or R2A media containing Congo Red for 48 hours prior to imaging. Each plate consisted of three replicates and experiments were repeated at least three times. (B) Swim agar plates were inoculated in the center of each plate and incubated for 24 hours prior to imaging. Average swimming ring diameter was determined from three individual experiments consisting of three biological replicates each. (*) indicate statistically significant differences (p ≤ 0.005) compared to the wild type strain and (^a^) represents statistically significant differences (p ≤ 0.005) compared to the strain expressing DGC2884, both of which were measured by the student’s t-test. (C) Indicated strains were grown under conditions conducive to biofilm formation on vinyl coverslips for 72 hours prior to staining with Crystal Violet. Average absorbance values at 550 nm for each sample are shown in graph. Data is representative of at least three independent experiments consisting of at least 3 biological replicates per sample each time. (*) indicate statistically significant differences (p < 0.005) as measured by the student’s t-test. (^a^) represents a statistically significant difference (p < 0.005) when compared to expression of DGC2884 as measured by the student’s t-test. (D) Pellicle formation assays were used to compare the percentages of cells in pellicles for each sample after a period of 72 hours. Each bar represents three biological replicates from a single experiment. Experiment was performed twice with similar results. (*) indicate statistically significant differences (p < 0.005) compared to the wild type strain and (a) represents a statistically significant difference (p < 0.05) compared to the strain expressing DGC2884, both of which were measured by the student’s t-test. (E) Fluorescence micrographs of representative cells co-expressing the Vc2 Spinach aptamer with the indicated constructs. Scale bar represents 5 µm. (F) Diguanylate cyclase enzyme activity is shown as a bar graph comparing the mean fluorescence intensity measured across 50 cells per sample. Error bars represent the S.E.M. values. (*) indicate statistically significant differences (p < 0.005) as measured by the student’s t-test. Data shown represents three separate experiments.

Since increased levels of c-di-GMP are typically associated with decreased motility (11, 43, 44), we next tested whether overexpression of these diguanylate cyclases affected motility using a swim plate agar assay. As expected, overexpression of DGC2884 resulted in impaired motility compared to the control strain, which was partially restored in the DGC2884 AADEF variant (Fig 2B). We found that, in comparison to strains overexpressing DGC2884, expression of DGC2884ΔTM resulted in partial restoration of motility behavior reminiscent of that observed for strains expressing the DGC2884 AADEF mutant (Fig 2B). Together, these data suggest that a fully functional DGC2884 is required to modulate motility.

Next, we examined whether overexpression of these diguanylate cyclases influenced biofilm formation (Fig 2C). While each of these strains showed formation of biofilms on vinyl coverslips, the most robust biofilms were formed during expression of the wild type DGC2884, which was reduced in the DGC2884 AADEF and DGC2884ΔTM mutants (Fig 2C), further indicating the importance of both an active enzymatic site and the N-terminal transmembrane domain.

We also tested the effect of overexpression of each diguanylate cyclase on pellicle formation and calculated the percentage of cells in pellicles and found that overexpression of DGC2884 resulted in significantly increased pellicle formation when compared to the empty vector control (p < 0.005, t-test) (Fig 2D). While expression of DGC2884 AADEF and DGC2884ΔTM also resulted in more pellicle formation than the control (significantly more by DGC2884ΔTM, p < 0.05, t-test), they produced significantly less pellicle than that of wild type cells expressing the full-length DGC2884 (p < 0.05, t-test) (Fig 2D).

Lastly, we assessed the enzyme activity of each diguanylate cyclase, including the DGC2884 AADEF mutant, using a Vc2-Spinach aptamer which acts as a c-di-GMP biosensor (45). For this assay, the full length DGC2884 diguanylate cyclase and each of the variants were expressed in *E. coli* BL21 DE3 Star cells and the fluorescence intensity of cells was measured as an indicator of c-di-GMP levels (Fig 2E, 2F). Indeed, we found that cells expressing DGC2884 had significantly higher fluorescence intensity compared to control cells, consistent with DGC2884 being an active diguanylate cyclase. Furthermore, cells expressing the DGC2884 AADEF variant were significantly less fluorescent than cells expressing DGC2884, suggesting that the AADEF mutation indeed affected enzyme activity (Fig 2E, 2F). We also found that expressing DGC2884ΔTM resulted in little to no activity (Fig 2E, 2F). To verify that the genes encoding these diguanylate cyclases were expressed in these cells, we examined transcript levels using RT-PCR (Fig S3). Taken together, results from each of these assays confirm that both enzyme activity and the N-terminal transmembrane are critical to the function of DGC2884.

### Domain architecture and role of transmembrane domain in localization pattern of DGC2884

To gain further insight into the function of DGC2884, we performed a simple Protter analysis using the amino acid sequence of DGC2884 (46) and found that the sequence for DGC2884 is predicted to have two transmembrane domains at its N-terminus that make up a CHASE8 domain, followed by the GGDEF domain (Fig 3A). The presence of CHASE domains within various proteins, including diguanylate cyclases, have been described as having different sensory capacities (47–50), though precisely what these domains sense is still unknown.

**Figure 3.**
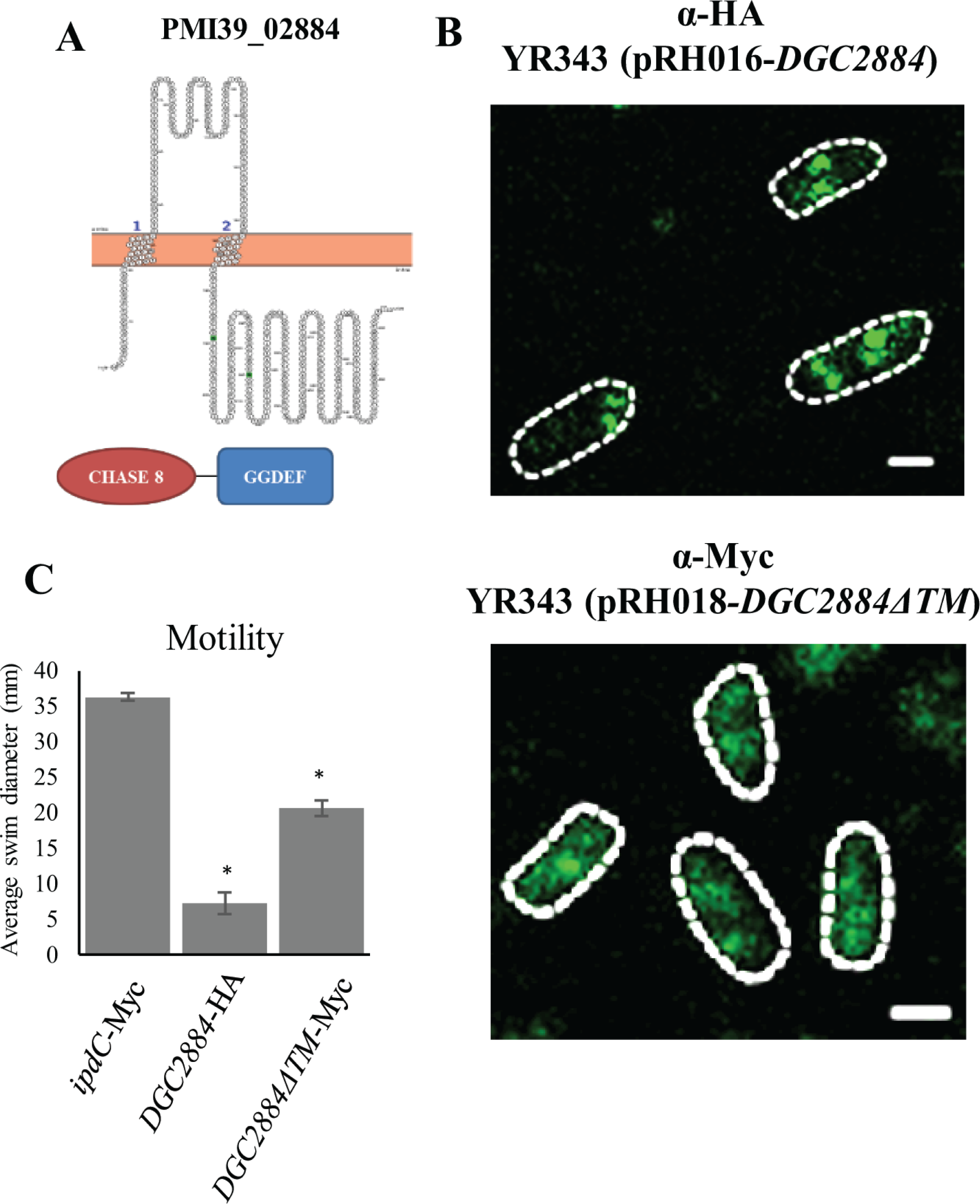
Localization of wild type DGC2884 and DGC2884ΔTM expressed in a wild type background using immunofluorescence assays. (A) Predicted topological structure of DGC2884 using Protter (top) and domain organization based on pfam analysis (bottom). (B) HA-tagged DGC2884 and Myc-tagged DGC2884ΔTM were detected using immunofluorescence assays and imaged using confocal microscopy. Individual cells are outlined in a white dashed line. Scale bars represent 1 µm. (C) Motility plate assays demonstrating functionality of tagged constructs, including an *ipdC* negative control. Data represent three biological replicates and at least two independent experiments. (*) indicate statistically significant differences (p < 0.05) as measured by the student’s t-test.

We next examined localization of wild type DGC2884 and DGC2884ΔTM in a wild type background by expressing it fused to either a 3HA or 13Myc tag (Fig 3B). These data show that DGC2884 was found to primarily localize in discrete foci at the cell pole or towards the mid-cell. In the absence of the N-terminal transmembrane domain, however, DGC2884 no longer localized as discrete foci, but rather the localization pattern became more diffuse with fewer visible foci (Fig 3B and Table 2). To verify that the tag did not alter the expression or function of these enzymes, we performed a motility assay (Fig 3C) and western blot (Fig S5) and observed that each construct was expressed and functional. As a control, we used a *Pantoea* YR343 strain carrying the same plasmid with *ipdC* inserted in place of the diguanylate cyclase gene where we do not expect to see any phenotypes associated with motility (27).

**TABLE 2.**
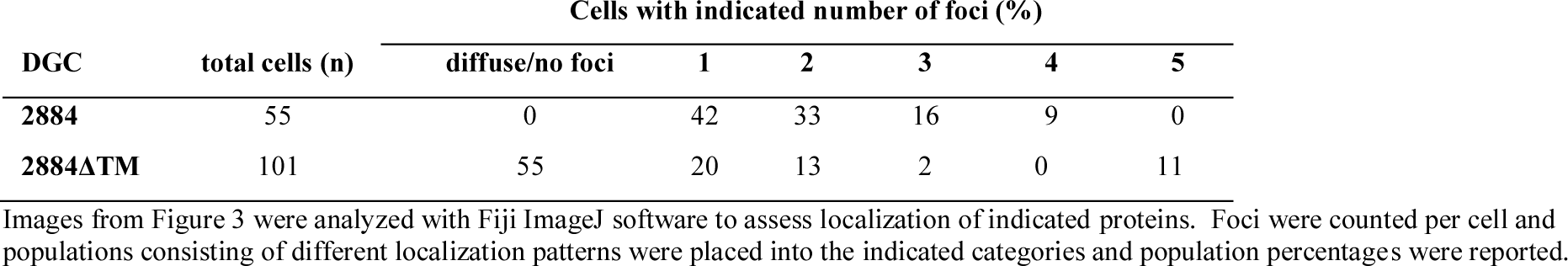
Quantification of GFP localization for tagged diguanylate cyclases.

### Identification of c-di-GMP responsive genes using transposon mutagenesis

Overexpression of DGC2884 resulted in a number of phenotypes (shown in Fig 2), including wrinkly colonies, increased Congo Red binding, increased pellicle and biofilm formation, and decreased motility, all of which suggest that DGC2884 is an active enzyme that influences c-di-GMP levels. Because expression of this diguanylate cyclase is influenced by plant association, we wanted to examine the molecular basis for the observed effects of DGC2884 expression on *Pantoea* behavior. To this end, we designed a transposon mutant screen to identify mutants that failed to respond to high levels of cyclic di-GMP resulting from DGC2884 overexpression, as determined by differences in Congo Red binding from that of the wild type strain expressing DGC2884. For this screen, we constructed a small transposon mutant library in *Pantoea* sp. YR343 (pSRK (Gm) –*DGC2884*) and screened for mutants of interest by plating the library on R2A plates containing Congo Red. Colonies that did not display the typical DGC2884 overexpression phenotype of wrinkled colonies and/or increased Congo Red binding were selected for further analyses. From this screen, we isolated 136 mutants that failed to respond to DGC2884 overexpression. Using a rescue cloning approach, we identified the location of the transposon insertion in each of these mutants, which resulted in a list of 61 genes, with some genes represented by multiple transposon mutants (Table 3). The top 5 COG categories represented among this set of genes include transcription (K), signal transduction (T), cell wall/membrane/envelope biogenesis (M), carbohydrate transport and metabolism (G), and intracellular trafficking/secretion/vesicular transport (U). We also identified 14 genes that were classified as either hypothetical proteins or that were not in a COG category.

**TABLE 3.**
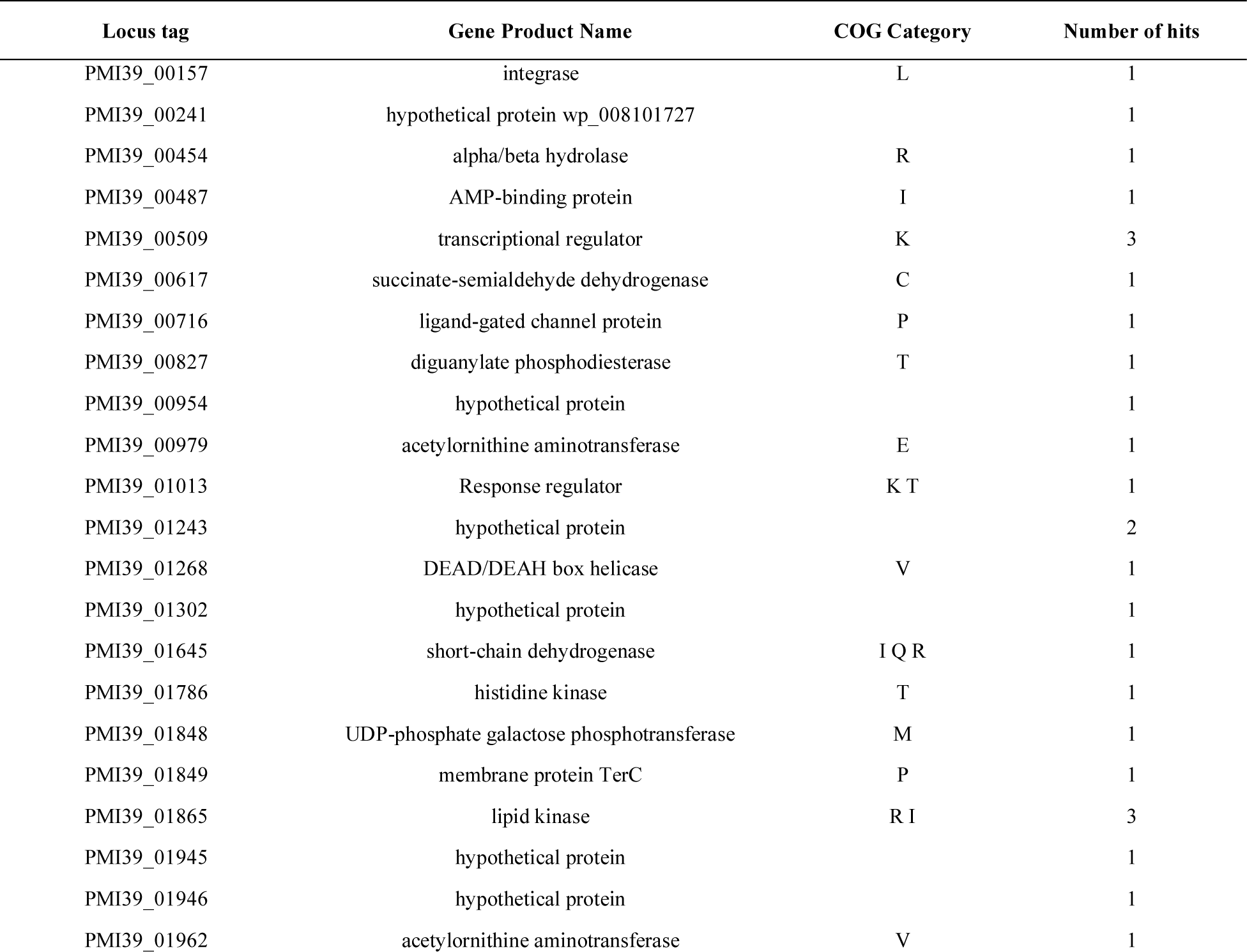
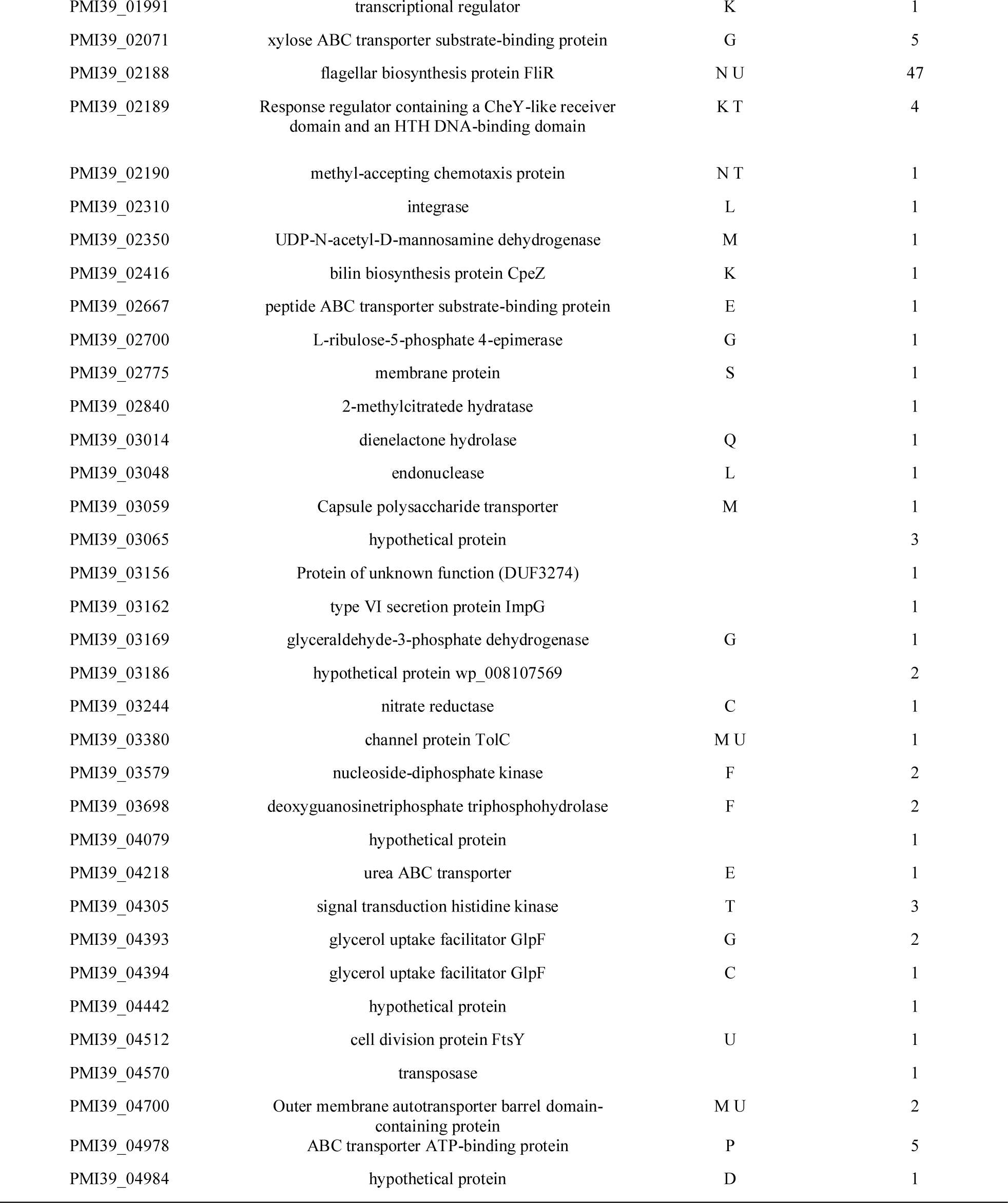
List of genes identified by transposon mutagenesis.

| Locus tag | Gene Product Name | COG Category | Number of hits |
| --- | --- | --- | --- |
| PMI39_00157 | integrase | L | 1 |
| PMI39_00241 | hypothetical protein wp_008101727 |  | 1 |
| PMI39_00454 | alpha/beta hydrolase | R | 1 |
| PMI39_00487 | AMP-binding protein | I | 1 |
| PMI39_00509 | transcriptional regulator | K | 3 |
| PMI39_00617 | succinate-semialdehyde dehydrogenase | C | 1 |
| PMI39_00716 | ligand-gated channel protein | P | 1 |
| PMI39_00827 | diguanylate phosphodiesterase | T | 1 |
| PMI39_00954 | hypothetical protein |  | 1 |
| PMI39_00979 | acetylornithine aminotransferase | E | 1 |
| PMI39_01013 | Response regulator | K T | 1 |
| PMI39_01243 | hypothetical protein |  | 2 |
| PMI39_01268 | DEAD/DEAH box helicase | V | 1 |
| PMI39_01302 | hypothetical protein |  | 1 |
| PMI39_01645 | short-chain dehydrogenase | I Q R | 1 |
| PMI39_01786 | histidine kinase | T | 1 |
| PMI39_01848 | UDP-phosphate galactose phosphotransferase | M | 1 |
| PMI39_01849 | membrane protein TerC | P | 1 |
| PMI39_01865 | lipid kinase | R I | 3 |
| PMI39_01945 | hypothetical protein |  | 1 |
| PMI39_01946 | hypothetical protein |  | 1 |
| PMI39_01962 | acetylornithine aminotransferase | V | 1 |
| PMI39_01991 | transcriptional regulator | K | 1 |
| PMI39_02071 | xylose ABC transporter substrate-binding protein | G | 5 |
| PMI39_02188 | flagellar biosynthesis protein FliR | N U | 47 |
| PMI39_02189 | Response regulator containing a CheY-like receiver domain and an HTH DNA-binding domain | K T | 4 |
| PMI39_02190 | methyl-accepting chemotaxis protein | N T | 1 |
| PMI39_02310 | integrase | L | 1 |
| PMI39_02350 | UDP-N-acetyl-D-mannosamine dehydrogenase | M | 1 |
| PMI39_02416 | bilin biosynthesis protein CpeZ | K | 1 |
| PMI39_02667 | peptide ABC transporter substrate-binding protein | E | 1 |
| PMI39_02700 | L-ribulose-5-phosphate 4-epimerase | G | 1 |
| PMI39_02775 | membrane protein | S | 1 |
| PMI39_02840 | 2-methylcitrate dehydratase |  | 1 |
| PMI39_03014 | dienelactone hydrolase | Q | 1 |
| PMI39_03048 | endonuclease | L | 1 |
| PMI39_03059 | Capsule polysaccharide transporter | M | 1 |
| PMI39_03065 | hypothetical protein |  | 3 |
| PMI39_03156 | Protein of unknown function (DUF3274) |  | 1 |
| PMI39_03162 | type VI secretion protein ImpG |  | 1 |
| PMI39_03169 | glyceraldehyde-3-phosphate dehydrogenase | G | 1 |
| PMI39_03186 | hypothetical protein wp_008107569 |  | 2 |
| PMI39_03244 | nitrate reductase | C | 1 |
| PMI39_03380 | channel protein TolC | M U | 1 |
| PMI39_03579 | nucleoside-diphosphate kinase | F | 2 |
| PMI39_03698 | deoxyguanosinetriphosphate triphosphohydrolase | F | 2 |
| PMI39_04079 | hypothetical protein |  | 1 |
| PMI39_04218 | urea ABC transporter | E | 1 |
| PMI39_04305 | signal transduction histidine kinase | T | 3 |
| PMI39_04393 | glycerol uptake facilitator GlpF | G | 2 |
| PMI39_04394 | glycerol uptake facilitator GlpF | C | 1 |
| PMI39_04442 | hypothetical protein |  | 1 |
| PMI39_04512 | cell division protein FtsY | U | 1 |
| PMI39_04570 | transposase |  | 1 |
| PMI39_04700 | Outer membrane autotransporter barrel domain-containing protein | M U | 2 |
| PMI39_04978 | ABC transporter ATP-binding protein | P | 5 |
| PMI39_04984 | hypothetical protein | D | 1 |

### Behavioral defects observed in selected mutants

Using the list of genes found in the genetic screen (Table 3), we selected eight different transposon mutants for further analyses, including a predicted UDP-galactose lipid carrier transferase (PMI39_01848; UDP::Tn5) and a predicted capsule polysaccharide transporter (PMI39_03059; CAP::Tn5), both of which have a predicted role in exopolysaccharide production. Because approximately one third of the identified transposon mutants were affected in *fliR* (PMI39_02188; FliR::Tn5), we included this mutant for further characterization as well. We also chose mutants predicted to be affected in Type VI secretion (PMI39_03162; Type VI::Tn5), in glycerol uptake (PMI39_04394; GlpF::Tn5), transport (PMI39_04218; ABC::Tn5), a nucleoside-diphosphate kinase (PMI39_03579; Ndk::Tn5), and one of the three hypothetical proteins (PMI39_03065; Hypo::Tn5). We began by curing each mutant of the DGC2884 expression plasmid (pSRK(gm)-*DGC2884*) in order to introduce an empty pSRK(gm) vector control prior to examining EPS production (by observing phenotypes on media with Congo Red) (Fig 4A, 4B). Next, we used the cured transposon mutants to observe pellicle formation (Fig 4C), and measure biofilm production with a crystal violet assay (Fig 4D). Compared to the wild type control, each mutant had a different growth phenotype on media with Congo Red, some of which were more noticeable on one media type over the other (Fig 4A, 4B). These phenotypes were further influenced based on whether the mutant expressed DGC2884 (pSRK (gm)-*DGC2884*) or an empty vector (pSRK (gm)). We next examined the effects of these mutations on pellicle formation and found that the UDP::Tn5, FliR::Tn5, and GlpF::Tn5 mutants produced significantly less pellicle than the wild type strain (Fig 4C). We also examined biofilms attached to vinyl coverslips and found that while some mutants appear to produce more biofilm, such as FliR::Tn5 and GlpF::Tn5, there were no statistically significant differences measured by quantifying Crystal Violet staining. Interestingly, we did find that the UDP::Tn5 and Ndk::Tn5 mutants produced significantly more biofilm than the wild type strain in this assay (Fig 4C).

**Figure 4.**
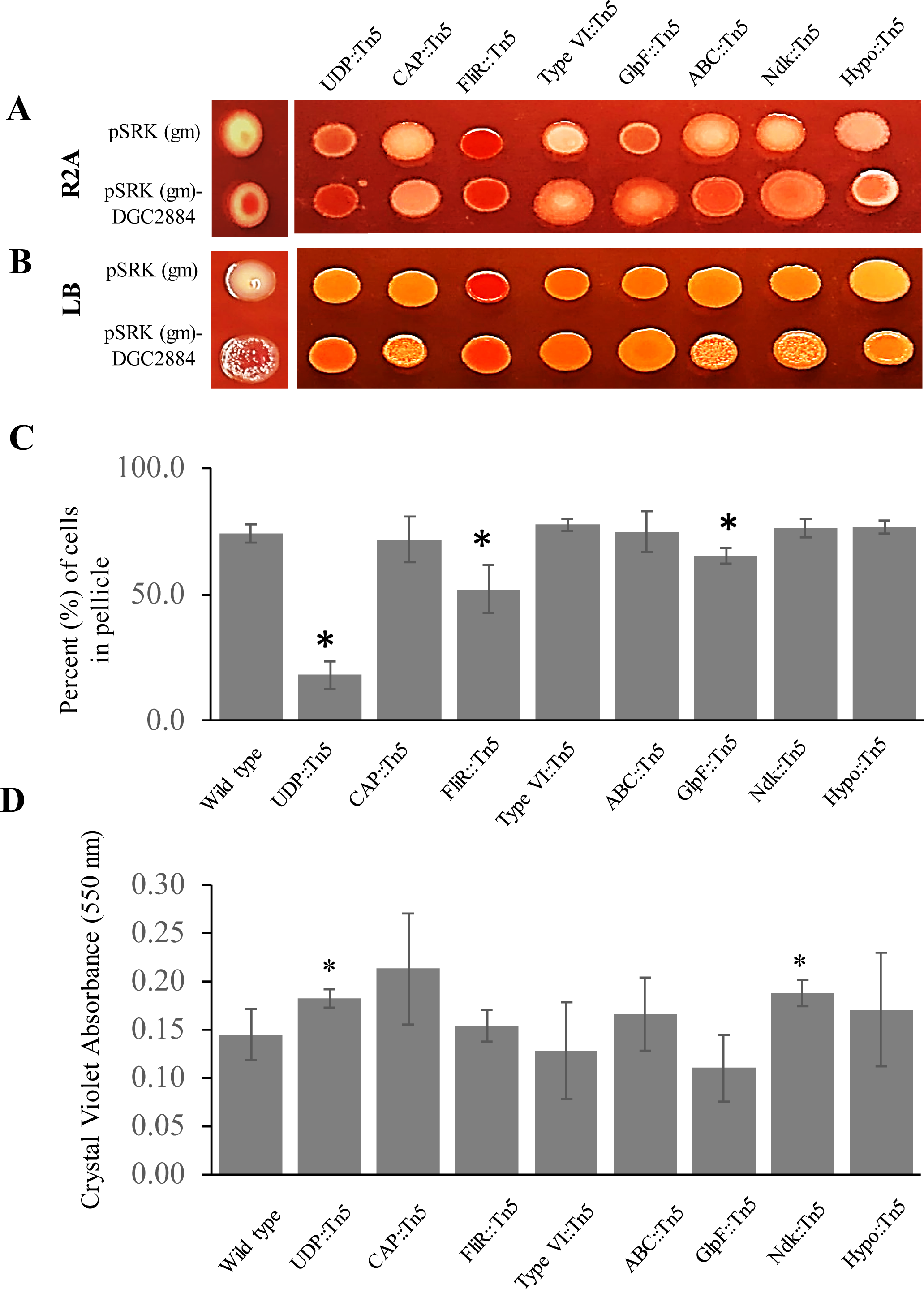
Characterization of behavioral defects among selected transposon mutants: UDP::Tn5 (PMI39_01848), CAP::Tn5 (PMI39_03059), FliR::Tn5 (PMI39_02188), TypeVI::Tn5 (PMI39_03162), GlpF::Tn5 (PMI39_04394), ABC::Tn5 (PMI39_04218), Ndk::Tn5 (PMI39_03579), and Hypo::Tn5 (PMI39_03065). Individual strains possessing either an empty pSRK-gm vector or pSRK(gm)-DGC2884 were spotted onto R2A (A) or LB (B) plates with Congo Red and incubated for 48 hours prior to imaging. (C) Each strain was grown under conditions conducive to pellicle formation for 72 hours. Graph indicates the average percentage of cells within the pellicles taken from three biological replicates. (*) indicate statistically significant differences (p < 0.005) when compared to the wild type strain using the student’s t-test. (D) Biofilm assays were performed using vinyl coverslips for 72 hours prior to staining with crystal violet. Bar graphs describe the average absorbance at 595 nm per sample as determined from two experiments, each with a set of three biological replicates. (*) indicates statistically significant differences (p < 0.005) when compared to the wild type strain using the student’s t-test.

While there were a variety of interesting behaviors observed for individual mutants in formation of biofilms and pellicles, these behaviors do not necessarily reflect what takes place in the rhizosphere during root colonization. We therefore chose to further characterize each of these mutants in colonization of *Populus* plant roots. For these studies, we examined colonization behavior of each mutant individually, and found that the UDP::Tn5 mutant showed significantly reduced colonization compared to the wild type strain, while the Type VI::Tn5, ABC::Tn5, and Ndk::Tn5 mutants showed a slight, though significant, increase in colonization (Fig 5A; statistically significant differences with p < 0.005, t-test). Comparisons of growth rates between transposon mutants and the wild type strain showed no significant differences for most strains, except for growth with UDP::Tn5 (Fig S4); however, based on growth curves, the maximum OD reached by wild type cells is approximately 0.67 (corresponding to a cell count of 5.36 x 10^8^ cells per mL), while the maximum OD for the UDP::Tn5 mutant is approximately 0.57 (corresponding to a cell count of 4.56 x 10^8^ cells per mL) which may contribute, in part, to the observed two orders of magnitude reduction in colonization.

**Figure 5.**
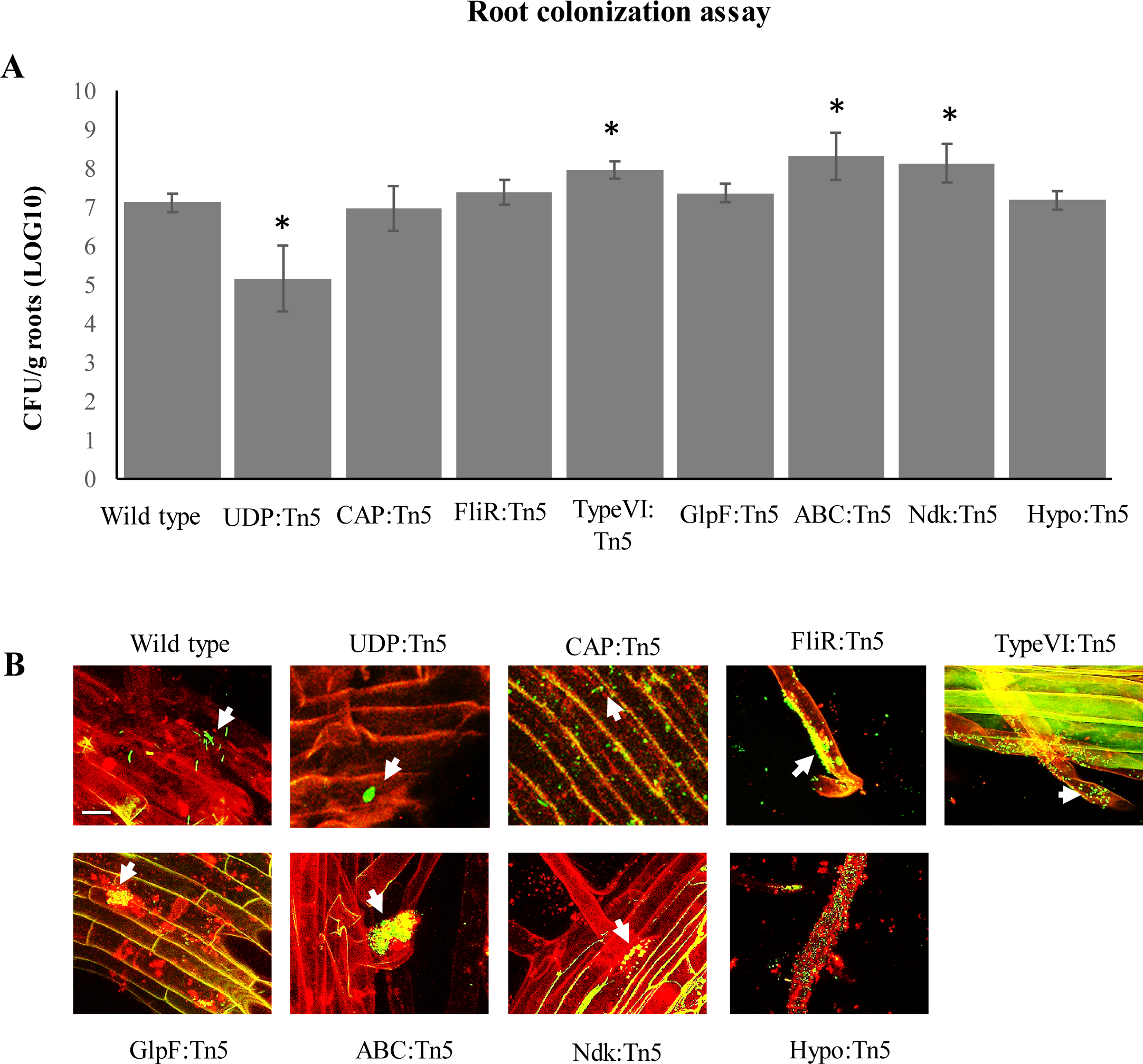
Effects of transposon mutations on root colonization patterns in *P. trichocarpa*. (A) Colonization of plant roots inoculated with wild type *Pantoea* YR343, UDP::Tn5, CAP::Tn5, FliR::Tn5, Type VI::Tn5, GlpF::Tn5, ABC::Tn5, Ndk::Tn5, or Hypo::Tn5 mutants and was measured after 3 weeks by counting CFUs relative to the total weight of each plant root. Error bars represent standard deviation over three to five different biological replicates per sample and (*) represent statistically significant differences (p ≤ 0.005, t-test). (B) Images of *P. trichocarpa* (red) one week after inoculation with the indicated bacterial strains (green). Scale bar represents 10 µm.

In addition to counting the overall number of microbes, we also wanted to determine how these mutant strains were distributed during colonization compared to wild type. For our studies, the wild type strain, the UDP::Tn5, and CAP::Tn5 mutant were each tagged with green fluorescent protein to facilitate imaging. The remaining mutants were observed by staining with Syto 9 dye (Fig 5B). In recent work by Noirot-Gros *et al*. (51), several patterns of root surface colonization in Aspen were described to facilitate comparisons, including no pattern (NP), long strips (LS), long patch microcolonies (LP), short patch microcolonies (SP), and high density bacterial coating (C). Using these descriptions, we observed that wild type *Pantoea* sp. YR343 exhibited a combination of long strips, long patch microcolonies, and small patches and was localized along the main root and root hair regions (Fig 5B). Consistent with its reduced counts, it was difficult to detect the UDP::Tn5 mutant on the root surface, although some small patches along the primary root surface were observed. The colonization pattern of CAP::Tn5 was very different from wild type, even though the overall level of colonization was similar (Fig 5B). Interestingly, the CAP::Tn5 mutant did not display a pattern of any kind, but was spread out over the root surface as individual cells and did not form patches like the wild type strain, possibly indicating a role of EPS in modulating the physical patterns of colonization along plant roots. Colonization by the FliR::Tn5, Type VI::Tn5, ABC::Tn5, and Hypo::Tn5 mutants consisted of long strips and small patches of cells and were found predominantly along the root hairs (Fig 5B). Both the Ndk::Tn5 and GlpF::Tn5 mutants exhibited primarily small patches of cells which appeared to be quite spread out along the main root, but near root hair regions (Fig 5B). Among these transposon mutants, the strains with the most noticeable differences in colonization patterns were CAP::Tn5, Ndk::Tn5, and GlpF::Tn5.

## Discussion

We describe here the identification of three diguanylate cyclases that were expressed consistently during colonization of plant roots. Although we only identified three diguanylate cyclase genes (out of 21 predicted genes) that are expressed in the presence of plants, it is possible that the nature of this assay may have excluded other potential genes of interest. For example, predicted promoter sites may not have been correct for some diguanylate cyclase genes, or some genes may have been expressed at earlier or later times during colonization. Further, some of the diguanylate cyclases that were not tested based on higher levels of expression (over 2.00) under normal growth conditions may have important roles during plant colonization. To date, we know of only one other diguanylate cyclase, Chp8 from *Pseudomonas syringae* pv. Tomato DC3000, that has been shown to be activated in the presence of plants and appears to be involved in reducing flagellin expression, increasing EPS production, and avoiding plant immune responses (52). Interestingly, the ability to suppress flagellar gene expression is a key strategy in avoiding plant immune responses since flagellin is a common pathogen-associated molecular pattern (PAMP) protein (53). While this appears to be a common strategy among plant pathogens for host invasion, there have been only a few reports describing the role of c-di-GMP in these processes (54).

We focused our characterization studies on DGC2884 which exhibited the most dramatic phenotypes when overexpressed, including modulating EPS production, motility, and biofilm formation. Perhaps one of the more intriguing attributes of DGC2884 is the N-terminal CHASE domain which appears to be necessary for localization and enzyme function. Furthermore, we found that the full-length DGC2884 localized as discrete foci within the cell, suggesting that this enzyme may influence local concentrations of c-di-GMP or may localize in order to yield specificity in its downstream effects. As expected, deletion of the N-terminal transmembrane domain impacted localization of DGC2884; in addition, given the behavioral defects observed in this mutant, these data together further support the important role of this transmembrane domain to the enzyme activity of DGC2884. As mentioned previously, the domain architecture and gene neighborhood of DGC2884 resembles that of YfiN from various bacterial species. Although these proteins are not entirely the same, some parallels can be drawn. For example, YfiN in *P. aeruginosa*, *E. coli*, and *S. enterica* have each been shown to impact production of small colony variants, swimming motility and EPS production, similar to DGC2884 (34, 35). Interestingly, YfiN has been shown to modulate production of Psl polysaccharides, whose operon possesses genes also found to regulate amylovoran biosynthesis in *Erwinia amylovora* (33, 34, 55). While the specific stimuli for each of these enzymes is not yet known, some studies suggest a reductive or osmotic stress may serve as an input to this system (33, 34, 56). This may imply that the chemical environment of the rhizosphere induces expression of this diguanylate cyclase; however, further experimentation is required to test this. The YfiN enzyme has been found in microbial isolates from many different environments, ranging from cystic fibrosis patients to the rhizosphere of *Populus,* raising intriguing questions about what types of environmental stimuli activate this enzyme and how that translates into downstream behaviors. Further studies of enzymes like DGC2884 may provide important insights into the functions of both medically-important and agriculturally important microbes, such as the closely related pathogens *Erwinia amylovora* or *Pantoea stewartii* (57, 58).

Transposon mutagenesis of *Pantoea* sp. YR343 (pSRK-*DGC2884*) resulted in 136 mutants that affected a total of 61 genes identified from a small-scale library consisting of approximately 5,000 different clones. Surprisingly, approximately one-third of these mutants were affected in a component of the flagellar export apparatus, *fliR*, and we are currently investigating the role of this protein in relation to DGC2884. Although we obtained a wide selection of mutants, the conditions of our mutagenesis assay likely did not reach saturation, yielding the possibility that there are other genes of interest that have not yet been identified. Among the transposon mutants described here, the phenotypes we observed for the UDP::Tn5, CAP::Tn5, GlpF::Tn5, Ndk::Tn5, and FliR::Tn5 mutants in biofilm formation and root colonization yielded some interesting insights into possible roles of these genes in the rhizosphere. The UDP::Tn5 and CAP::Tn5 mutants each have predicted roles in exopolysaccharide biosynthesis and transport (59). The GlpF::Tn5 mutant affects an aquaglyceroporin that is involved in water and glycerol uptake and may be involved in osmoregulation (60). The Ndk::Tn5 mutant affects a nucleoside diphosphate kinase that has been shown to have a role in regulating cell growth and signaling, as well as cell surface polysaccharides in *P. aeruginosa* and *Mycobacterium tuberculosis* (61). Lastly, FliR is a member of the flagellar export apparatus(62–64).

Exopolysaccharides are known to play an important role in biofilm formation and root colonization (7, 9, 65). Many *Pantoea* species are associated with plants, some as pathogens while others appear to be beneficial (66). Two closely related species, *P. stewartii* and *E. amylovora*, have been shown to produce specific types of EPS, known as stewartan and amylovoran, that are integral to their function as pathogens (58, 67–69). Stewartan and amylovoran are involved in clogging the flow of xylem in plant tissues and causing Stewart’s wilt disease in corn and fire blight in various fruit trees, such as pears and apples (68, 70). The composition of EPS is similar between stewartan and amylovoran and consists primarily of galactose, glucose, and glucuronic acid (71–73). While the composition and structure of EPS produced by *Pantoea* sp. YR343 has not yet been characterized, the genome does encode a large operon that is homologous to the operons found to be responsible for production of stewartan and amylovoran in *P. stewartii* and *E. amylovora*, respectively. The described UDP-galactose lipid carrier transferase (UDP) gene, for which we have a mini-Tn5 insertion, is the first gene in this operon and it is likely that disruption of this gene affects the entire operon. Interestingly, *Pantoea* sp. YR343 has two genes with similarity to the UDP-galactose lipid carrier transferase (PMI39_01848 and PMI39_04793) and analysis of closely related species has shown that the presence of more than one UDP-galactose lipid carrier transferase is common. The other UDP gene (PMI39_04793) is annotated as WbaP and is most likely involved in lipopolysaccharide biosynthesis. To date, there has been extensive research done to describe lipopolysaccharide biosynthesis, including the role of WbaP in that process (for some reviews, see (74, 75)). While studies of this EPS have focused primarily on its role in pathogenesis, we have not found any evidence to suggest that *Pantoea* sp. YR343 is pathogenic under the conditions tested (25); rather, we have found that *Pantoea* sp. YR343 is a robust root colonizer and mutations affecting EPS result in a significant reduction in root colonization. We show here that the UDP::Tn5 mutant shows little observable root colonization, while the CAP::Tn5 mutant colonizes well, but in a pattern that differs significantly from the wild type. Interestingly, there were no indications of a colonization defect in the CAP::Tn5 mutant based on cell number per gram root; however, imaging studies indicated that the CAP::Tn5 mutant does not form patches of cell aggregates on the root surface, suggesting differences in the amount or composition of EPS in this mutant. Further studies into the composition and structure of the EPS from *Pantoea* sp. YR343 may yield insights into its function during plant association. Furthermore, as mentioned previously, it has been found that the YfiN enzyme can regulate production of the Psl polysaccharide in *P. aeruginosa*, suggesting a possible linkage to activation of EPS production by DGC2884 in *Pantoea* sp. YR343.

Lastly, we did observe an increase in root colonization by the Type VI::Tn5, ABC::Tn5, and Ndk::Tn5 mutants, although the increases were slight. Interestingly, the patterns of colonization in the Ndk::Tn5 mutant were mostly in the form of small patches along the root surface, while colonization by the TypeVI::Tn5 and ABC::Tn5 mutants were observed as both small patches and long strips along the root hairs. It is interesting to speculate that some of these genes may be involved in regulating colonization location, as well as pattern formation during root colonization. For example, the Type VI secretion system has been shown to be fairly widespread across plant-associated proteobacteria where the Type VI system is believed to promote fitness and competition within the rhizosphere during root colonization (76). Perhaps without this system, the cells are unable to colonize the older tissue along the primary plant root, but can better colonize the softer root hairs. More studies will be required to understand how these different gene products are involved in regulating colonization behaviors. Techniques involving imaging of spatial colonization patterns along plant roots, in addition to quantification by cell counts, can provide additional information about how these bacteria behave in the environment and how these mutations affect that behavior.

In summary, we have shown that growth in the presence of a plant host results in expression of genes encoding the diguanylate cyclases, DGC2884, DGC3006, and DGC3134 in *Pantoea* sp YR343. While similarities between YfiN and DGC2884 suggest environmental stresses as a stimulus, further characterization is needed to understand what triggers expression of these c-di-GMP signaling pathways in the rhizosphere. While there were 61 genes identified through transposon mutagenesis, we have only just begun to characterize how those gene products influence root colonization behavior and how they function in that process. Finally, the finding that at least three diguanylate cyclases were expressed on plant roots suggests how important c-di-GMP signaling is for the process of root colonization by *Pantoea* sp. YR343. Characterizing the coordination of these three diguanylate cyclases in the process of root colonization by *Pantoea* sp. YR343 will enhance our understanding of c-di-GMP regulation across bacteria and yield important insights into the roles of multiple diguanylate cyclases in coordinating these behaviors.

## Materials and Methods

### Bacterial strains and growth conditions

Table S1 describes the bacterial strains and plasmids used throughout this study. *E. coli* strains were grown in Luria Broth (LB) media (10 g Tryptone, 5 g Yeast Extract, and 10 g NaCl per 1 liter) with shaking at 37°C. *Pantoea* strains were grown in either R2A broth (TEKnova, Inc.), LB, TY (10 g tryptone and 5 g yeast extract per 1 liter), or M9 media with 0.4% glucose and grown at 28°C. Growth curve assays were performed using 96-well plates in a Biotek Cytation 5 plate reader. Motility assays were performed on low agar plates prepared by adding 0.3% agar to LB media. Congo Red plates (for EPS analysis) were prepared by adding Congo Red to R2A or LB media at a concentration of 40 µg ml^-1^.

### Construction of promoter-reporter plasmids

To generate reporter constructs, we analyzed the genomic sequences upstream of the predicted ATG start sites of each diguanylate cyclase in order to identify putative promoters using BPROM (31). We then cloned 200 bp regions encoding the predicted prometers for use in the reporter construct, pPROBE-NT (30). Primers used to amplify each of these promoter regions are listed in Table S2. Final constructs were verified using restriction digests, prior to introduction into *Pantoea* sp. YR343 using electroporation.

### Construction of diguanylate cyclase expression vectors

DGC2884 and DGC2884ΔTM were amplified by PCR from genomic DNA using the primers listed in Table S2. PCR products were digested with BamHI and HindIII before ligation into pSRK-Km or pSRK-Gm(77). The DGC2884 AADEF mutant was generated using a QuikChange Site-directed mutagenesis kit, with the cloned wild type DGC2884 as a template, and the resulting construct was verified by sequencing. Each construct was verified before transforming into electrocompetent wild type *Pantoea* sp. YR343, as described previously (25). Each overexpression strain was maintained with either 50 µg mL^-1^ kanamycin (pSRK-Km) or 10 µg mL^-1^ gentamycin (pSRK-Gm) and induction was performed by adding 1 mM isopropyl β-D-1-thiogalactopyranoside (IPTG).

Tagged diguanylate cyclases were generated by PCR amplification of each diguanylate cyclase followed by cloning into pENTR D-TOPO (ThermoFisher Scientific) and then transferred to either pRH016 (HA tag) or pRH018 (Myc tag) (78). Final constructs containing HA or Myc tags were then introduced into *Pantoea* YR343 via electroporation as described previously (25).

### Biofilm formation assays

We tested biofilm formation on vinyl coverslips using cultures grown in M9 minimal media supplemented with 0.4% glucose. Coverslips were sterilized by soaking in 100% bleach for 20 minutes and then rinsed twice in sterile water before placing in a sterile 12-well tissue culture dish. Sterility was tested using sterile M9 media with 0.4% glucose with sterilized coverslips. Cultures were grown in M9 minimal media overnight, then diluted 1:100 into fresh M9 media with 0.4% glucose (1 mM IPTG was added for strains with pSRK). Diluted cultures (1 mL) were then placed into a 12-well tissue culture dish (three replicates per strain) along with a sterilized vinyl coverslip placed at an angle. A breathable cover was placed over the 12-well tissue culture dish and was placed at 28°C for 72 hours. Coverslips were then removed and rinsed in water prior to staining with crystal violet as described previously (25). Expression using GFP reporter strains was measured by rinsing coverslips and imaging large sections of biofilms over a minimum of three separate fields of view. Details of image analysis are described below.

### Pellicle formation assays

Pellicle assays were performed as described previously (25). In order to quantify the percentage of cells within each pellicle, we collected 100 µL of cells from the non-pellicle portion of each culture, then used a glass homogenizer to disperse the pellicle with the remaining culture. From the homogenized culture, we collected 100 µL of cells. The OD (600 nm) was measured for the homogenized culture, as well as the non-pellicle portion, in order to calculate the percentage of cells that were in the pellicle. There were three biological replicates measured per experiment. Expression using GFP reporter strains was measured by placing pellicle cultures in a 96-well dish and normalizing fluorescence measurements to cell density (OD at 600 nm) as measured using the Biotek Cytation 5 plate reader.

### Expression of diguanylate cyclases and enzyme assays using Vc2-Spinach

We obtained the pET31b-Vc2 Spinach vector as a gift from Dr. Ming Hammond and used *E. coli* BL21 DE3 Star cells to co-express the Vc2-Spinach tRNA (from pET31b-Vc2 Spinach) with each diguanylate cyclase construct (pSRK (Km) DGC2884, pSRK (Km) DGC2884 AADEF, pSRK (Km) and DGC2884ΔTM) individually as described previously (45). We verified expression of each of these constructs by RT-PCR using protocols described below (see *Expression analysis*). Measurement of c-di-GMP using the Vc2-Spinach aptamer was performed as described previously (45).

### Expression analysis

In order to ensure expression of diguanylate cyclase genes, we performed RT-PCR using primer sets for each diguanylate cyclase (primers listed in Table S2). RNA extraction was performed using the Qiagen RNeasy Mini Kit according to manufacturer’s protocols. The SuperScript IV RT-PCR system (ThermoFisher Scientific) was used for generating cDNA according to manufacturer’s procedures and final PCR reactions were performed according to standard protocols.

### Immunolocalization and Western blotting

Immunolocalization was performed on *Pantoea* sp. YR343 (pRH016-*DGC2884*) and YR343 (pRH018-*DGC2884ΔTM*) using mouse polyclonal antibodies against HA (ab16918 from Abcam) or Myc (13-2500 from Invitrogen). Approximately 3 mL of cell culture grown to a low cell density was collected, washed in PBS, and fixed in ice cold methanol for 1 hour at −20°C. Afterwards, cells were placed onto poly-L-lysine coated coverslips and allowed to dry before lysing with 2 mg mL^-1^ lysozyme solution in GTE buffer (50 mM glucose, 20 mM Tris-HCl pH 8.0, 10 mM EDTA) for 10 minutes, followed by incubation overnight at 4°C in a blocking solution consisting of 1% non-fat dry milk in PBS, pH 7.0. After washing twice with PBS, the antibody solution consisting of 1% non-fat dry milk in PBS with either 1:250 dilution of anti-HA or 1:125 dilution of anti-Myc antibody was added and incubated for 2 hrs at room temperature. The coverslips were rinsed in PBS and then incubated with Alexa Fluor 488 goat anti-mouse IgG at a 1:500 dilution in 1% non-fat dry milk in PBS for 2 hours at room temperature. Coverslips were washed in PBS, mounted on slides and imaged using a Zeiss LSM710 confocal microscope.

Lysates were prepared for western blotting by collecting cell pellets from cultures grown overnight, then lysed by sonication. The crude lysate was centrifuged and supernatants were used for SDS-PAGE gels. Western blotting was performed according to standard protocols using the same mouse monoclonal antibodies used for immunolocalization.

### Transposon mutagenesis and rescue cloning

Biparental mating was used to introduce the plasmid pRL27, encoding a mini-Tn5 transposon, into *Pantoea* sp. YR343 (DGC2884 pSRK-Gm) essentially as described previously, but on a smaller scale (79). Removal of *E. coli* strain EA145 was performed by growing *Pantoea* in the presence of kanamycin (50 µg mL^-1^) and gentamycin (10 µg mL^-1^). Screening of the transposon library was performed by plating the library onto LB plates containing Congo Red (40 µg ml^-1^), 1 mM IPTG, kanamycin (50 µg mL^-1^) and gentamycin (10 µg mL^-1^). Colonies differing in appearance from the parental strain were isolated for further characterization and for sequencing.

In order to identify the location of transposon insertions, we used a cloning approach described previously (79). Basically, we isolated genomic DNA from each mutant using the Promega Wizard Genomic DNA Extraction Kit, digested with a single restriction enzyme (most often used EcoRI, but sometimes used BamHI, PstI, SalI, SacII, and SphI) that does not cut within the transposon, ligated the DNA into plasmids, transformed these plasmids into *E. coli* PIR1 cells (ThermoFisher Scientific) and then plated onto selective plates containing 50 µg mL^-1^ kanamycin (transposon sequence contained a kanamycin resistance marker). Colonies were picked and plasmid DNA was isolated using the QIAprep Spin Miniprep Kit (Qiagen) and plasmids were sequenced at the Molecular Biology Resource Facility at the University of Tennessee, Knoxville. We sequenced each plasmid from the transposon outwards using the following primers, tpnRL17-1 and tpnRL13-1 (79). All resulting sequences were analyzed using BlastX from NCBI in order to identify the region of DNA flanking each transposon.

Individual transposon mutants were grown three to four times sequentially on rich media without selection in order to remove the pSRK (Gm)-*DGC2884* plasmid. Removal of the plasmid was verified by growth on kanamycin at 50 µg mL^-1^, but not on gentamycin at 10 µg mL^-1^.

### Construction of fluorescent strains

We generated fluorescent strains that were also resistant to gentamycin by integrating either GFP or mCherry into the chromosome of *Pantoea* sp. YR343 using the pBT270 and pBT277 plasmids which use the Tn7 transposon system for chromosomal insertions (gift from B.S. Tseng and(80). Colonies with chromosomally inserted GFP or mCherry were selected on R2A agar containing gentamycin at 10 µg mL^-1^.

### Plant growth conditions and inoculation

Wheat seeds were grown in special growth chambers (Advanced Science Tools, LLC – http://advancedsciencetools.com/index.html) which allow for visualization of plant roots without sacrificing the plants. Wheat seeds were surface-sterilized, as performed previously (25) and placed into the chamber filled with sterile soil. Once seeds were germinated and had both stem and roots, plants were inoculated with *Pantoea* sp. YR343, as described previously (25). Plants were incubated with *Pantoea* sp. YR343 for 7 days prior to visualization using confocal microscopy.

Colonization of *Populus trichocarpa* BESC819 was performed as described previously (25). Five plants were used per treatment (with approximately 10^7^ cells per plant) and incubations were for three weeks prior to harvesting and counting. Visualization of non-fluorescently labelled cells was performed by staining with Syto 9, as described previously (25).

### Confocal fluorescence microscopy and image analysis

Biofilms were imaged for promoter expression analysis using a Biotek Cytation 5 plate reader. Images were taken from at least three separate fields-of-view per sample. In order to quantify fluorescence, we drew nine square regions of interest per image using Fiji ImageJ and measured fluorescence intensity per square for all images. The average fluorescence intensity per square micrometer, as well as the S.E.M. values were calculated for each sample and then normalized against the pPROBE empty vector control.

Confocal fluorescence microscopy was performed using a Zeiss LSM710 confocal laser scanning microscope with a Plan-Apochromat 63x/1.40 oil immersion objective (Carl Zeiss Microimaging, Thornwood, NY). Images were processed using Zen2012 software (Zeiss). Cell fluorescence intensity measurements were performed using Fiji ImageJ for assays with promoter-reporter fusions for DGCs and for the Vc2 Spinach aptamer following the protocol described by Kellenberger, et al (45). Briefly, images were initially collected using the same parameters and then collectively processed so that brightness and contrast was adjusted and normalized across the entire set of images used for analysis. Using brightfield images, individual regions-of-interest were drawn for a minimum of 50 cells, then used to measure fluorescence in corresponding fluorescent images.

## Acknowledgements

We would like to thank Dr. B. S. Tseng (University of Washington) for sharing plasmids pBT270 and pBT277, as well as Dr. Alison Buchan (University of Tennessee, Knoxville) for sharing the pRL27 plasmid, and finally, Dr. Ming Hammond (University of Utah) for sharing pET31b-Vc2 Spinach. This research was sponsored by the Genomic Science Program, U.S. Department of Energy, Office of Science, Biological and Environmental Research, as part of the Plant Microbe Interfaces Scientific Focus Area (http://pmi.ornl.gov). Oak Ridge National Laboratory is managed by UT-Battelle LLC, for the U.S. Department of Energy under contract DE-AC05-00OR22725.

## Supporting Information Captions

Figure S1. Clustal Omega multiple sequence alignment of *Pantoea* sp. YR343 DGC2884, *Pseudomonas aeruginosa* PA01 TpbB, and *Escherichia coli* MG1655 DgcN (36).

Figure S2. Growth curves of wild type (pSRK-Km) and the indicated DGC overexpressing strains in minimal media (A) and LB media (B). Error bars represent the standard deviation from three independent cultures.

Figure S3. Expression of individual diguanylate cyclases using RT-PCR. Image shown is representative of a minimum of 3 replicates.

Figure S4. Growth curves of wild type *Pantoea* sp. YR343 and indicated transposon mutants in minimal media (A) and in LB media (B). Error bars represent the standard deviation from three independent cultures.

Figure S5. Western blot showing expression of tagged full length DGC2884 and DGC2884ΔTM. Weights of markers are indicated on the left and arrows point to bands that represent the indicated protein.

