## Supplemental Data for "Identification of gene products involved in plant colonization by *Pantoea* sp. YR343 using a diguanylate cyclase expressed in the presence of plants"

### Supplementary Tables

TABLE S1. Strains and Plasmids used in this study

| Strain or plasmid | Genotype or relevant characteristics | Reference or source |
|---|---|---|
| Plasmids | | |
| pET DEST42 | Expression vector | Invitrogen |
| pDEST14 | Cloning vector | Invitrogen |
| pET DEST14-42 | Expression vector with C-terminal V5 and His tags | this work |
| pET31b-Vc2 Spinach | Expression vector with Vc2-Spinach aptamer | (42) |
| pPROBE-NT | Cloning vector with promoter-less GFP (Km | (73) |
| pSRK-Km | Expression vector, (Km) | (74) |
| pSRK-Gm | Expression vector, (Gm*) | (74) |
| pRL27 | Tn5-RL27 (Km$^R$-*ori*R6K) delivery vector | (76) |
| pRH016 | pBBR1 gateway expression vector with C-terminal 3HA tag, (Cm*) | (75) |
| pRH018 | pBBR1 gateway expression vector with C-terminal 13MYC tag, (Cm) | (75) |
| pPROBE-*DGC0366* | pPROBE containing promoter region of *DGC0366* | this work |
| pPROBE-*DGC0751* | pPROBE containing promoter region of *DGC0751* | this work |
| pPROBE-*DGC0995* | pPROBE containing promoter region of *DGC0995* | this work |
| pPROBE-*DGC1008* | pPROBE containing promoter region of *DGC1008* | this work |
| pPROBE-*DGC1023* | pPROBE containing promoter region of *DGC1023* | this work |
| pPROBE-*DGC1024* | pPROBE containing promoter region of *DGC1024* | this work |
| pPROBE-*DGC1089* | pPROBE containing promoter region of *DGC1089* | this work |
| pPROBE-*DGC1854* | pPROBE containing promoter region of *DGC1854* | this work |
| pPROBE-*DGC2196* | pPROBE containing promoter region of *DGC2196* | this work |
| pPROBE-*DGC2242* | pPROBE containing promoter region of *DGC2242* | this work |
| pPROBE-*DGC2465* | pPROBE containing promoter region of *DGC2465* | this work |
| pPROBE-*DGC2697* | pPROBE containing promoter region of *DGC2697* | this work |
| pPROBE-*DGC2884* | pPROBE containing promoter region of *DGC2884* | this work |
| pPROBE-*DGC3006* | pPROBE containing promoter region of *DGC3006* | this work |
| pPROBE-*DGC3134* | pPROBE containing promoter region of *DGC3134* | this work |
| pPROBE-*DGC3217* | pPROBE containing promoter region of *DGC3217* | this work |
| pPROBE-*DGC3247* | pPROBE containing promoter region of *DGC3247* | this work |
| pPROBE-*DGC3482* | pPROBE containing promoter region of *DGC3482* | this work |

| | | |
|---|---|---|
| pPROBE-*DGC3621* | pPROBE containing promoter region of *DGC3621* | this work |
| pPROBE-*DGC4070* | pPROBE containing promoter region of *DGC4070* | this work |
| pSRK (Km)-*DGC2884* | pSRK-Km containing full gene sequence of *DGC2884* | this work |
| pSRK (Km)-*DGC2884 AADEF* | pSRK-Km containing gene sequence of *DGC2884 AADEF* | this work |
| pSRK (Km)-*DGC2884ΔTM* | pSRK-Km containing full gene sequence of *DGC2884ΔTM* | this work |
| pRH016-*DGC2884* | pRH016 containing *DGC2884* with C-terminal 3HA tag | this work |
| pRH016-*DGC2884ΔTM* | pRH016 containing *DGC2884ΔTM* with C-terminal 3HA tag | this work |
| pRH018-*DGC2884ΔTM* | pRH018 containing *DGC2884ΔTM* with C-terminal 13MYC tag | this work |
| pBT270 | pUC18-miniTn7T2-PA1/04/03-GFP (Ap*, Gm) | gift from Dr. B. S. Tseng |
| pBT277 | pUC18-miniTn7T2-PA1/04/03-mCherry (Ap, Gm) | (78) |
| Strains | | |
| *Escherichia co* | | |
| TOP10 | General cloning strain | ThermoFisher Scientific |
| EA145 | Mating strain containing pRL27, DAP auxotroph | gift from Dr. A. Buchan |
| BL21 DE3 Star | Protein expression strain | ThermoFisher Scientific |
| *Pantoea* sp. | | |
| YR343 | wild type strain | |
| YR343 (pPROBE-*DGC0366*) | YR343 containing pPROBE-*DGC0366* | this work |
| YR343 (pPROBE-*DGC0751*) | YR343 containing pPROBE-*DGC0751* | this work |
| YR343 (pPROBE-*DGC0995*) | YR343 containing pPROBE-*DGC0995* | this work |
| YR343 (pPROBE-*DGC1008*) | YR343 containing pPROBE-*DGC1008* | this work |
| YR343 (pPROBE-*DGC1023*) | YR343 containing pPROBE-*DGC1023* | this work |
| YR343 (pPROBE-*DGC1024*) | YR343 containing pPROBE-*DGC1024* | this work |
| YR343 (pPROBE-*DGC1089*) | YR343 containing pPROBE-*DGC1089* | this work |
| YR343 (pPROBE-*DGC1854*) | YR343 containing pPROBE-*DGC1854* | this work |
| YR343 (pPROBE-*DGC2196*) | YR343 containing pPROBE-*DGC2196* | this work |
| YR343 (pPROBE-*DGC2242*) | YR343 containing pPROBE-*DGC2242* | this work |
| YR343 (pPROBE-*DGC2465*) | YR343 containing pPROBE-*DGC2465* | this work |
| YR343 (pPROBE-*DGC2697*) | YR343 containing pPROBE-*DGC2697* | this work |
| YR343 (pPROBE-*DGC2884*) | YR343 containing pPROBE-*DGC2884* | this work |
| YR343 (pPROBE-*DGC3006*) | YR343 containing pPROBE-*DGC3006* | this work |
| YR343 (pPROBE-*DGC3134*) | YR343 containing pPROBE-*DGC3134* | this work |
| YR343 (pPROBE-*DGC3217*) | YR343 containing pPROBE-*DGC3217* | this work |
| YR343 (pPROBE-*DGC3247*) | YR343 containing pPROBE-*DGC3247* | this work |
| YR343 (pPROBE-*DGC3482*) | YR343 containing pPROBE-*DGC3482* | this work |

| | | |
|---|---|---|
| YR343 (pPROBE-*DGC3621*) | YR343 containing pPROBE-*DGC3621* | this work |
| YR343 (pPROBE-*DGC4070*) | YR343 containing pPROBE-*DGC4070* | this work |
| YR343 (pSRK-Km) | YR343 containing pSRK-Km | this work |
| YR343 (pSRK-*DGC2884*) | YR343 containing pSRK-*DGC2884* (Km) | this work |
| YR343 (pSRK-*DGC2884 AADEF*) | YR343 containing pSRK-*DGC2884* AADEF (Km) | this work |
| YR343 (pSRK-*DGC2884ΔTM*) | YR343 containing pSRK-*DGC2884ΔTM* (Km) | this work |
| YR343 (pSRK(Gm)-*DGC2884*) | YR343 containing pSRK(Gm)-*DGC2884* (Gm) | this work |
| BL21 DE3 Star (pET31b-Vc2 Spinach + pSRK-Km) | BL21 DE3 Star containing pET31b-Vc2 Spinach (Cb) and pSRK-Km | this work |
| BL21 DE3 Star (pET31b-Vc2 Spinach + pSRK-*DGC2884*) | BL21 DE3 Star containing pET31b-Vc2 Spinach (Cb) and pSRK-*DGC2884* (Km) | this work |
| BL21 DE3 Star (pET31b-Vc2 Spinach + pSRK-*DGC2884ΔTM*) | BL21 DE3 Star containing pET31b-Vc2 Spinach (Cb) and pSRK-*DGC2884ΔTM* (Km) | this work |
| BL21 DE3 Star (pET31b-Vc2 Spinach + *pSRK-DGC2884 AADEF*) | BL21 DE3 Star containing pET31b-Vc2 Spinach (Cb) and pSRK-*DGC2884 AADEF*(Km) | this work |
| YR343 (pRH016-*DGC2884*) | YR343 containing pRH016-*DGC2884* (Cm) | this work |
| YR343 (pRH018-*DGC2884ΔTM*) | YR343 containing pRH018-*DGC2884ΔTM* (Cm) | this work |
| YR343 (pRH018-*ipdC*) | YR343 containing pRH018-*ipdC* (Cm) | Morrell-Falvey Lab |
| YR343:GFP | YR343 with GFP integrated chromosomally via pBT270 | this work |
| CAP::Tn5 | YR343 mutant with a transposon insertion in PMI39_03059 | this work |
| CAP:mCherry | CAP::Tn5 with mCherry integrated chromosomally via pBT277 | this work |
| UDP::Tn5 | YR343 mutant with a transposon insertion in PMI39_01848 | this work |
| UDP:mCherry | UDP::Tn5 with mCherry integrated chromosomally via pBT277 | this work |
| FliR::Tn5 | YR343 mutant with a transposon insertion in PMI39_02188 | this work |
| TypeVI::Tn5 | YR343 mutant with a transposon insertion in PMI39_03162 | this work |
| GlpF::Tn5 | YR343 mutant with a transposon insertion in PMI39_04394 | this work |
| ABC::Tn5 | YR343 mutant with a transposon insertion in PMI39_04218 | this work |
| Ndk::Tn5 | YR343 mutant with a transposon insertion in PMI39_03579 | this work |
| Hypo::Tn5 | YR343 mutant with a transposon insertion in PMI39_03065 | this work |

*Antibiotic resistance: Km, kanamycin; Gm, gentamycin; Ap, ampicillin; Cm, chloramphenicol

### TABLE S2. Primers used in this study

| Primer Name | Primer Sequence (5’ -> 3’) |
| --- | --- |
| DGC0366 prom-For | CGCGGATCCCCGTAGCAAAGTCAGGCC |
| DGC0366 prom-Rev | CCGGAATTCCCCTGTGCCTGCATCACT |
| DGC0751 prom-For | CGCGGATCCCGATGTGACTACCGAATG |
| DGC0751 prom-Rev | CCGGAATTCGTTGTCCTCTGTTTTCAA |
| DGC0995 prom-For | CGCGGATCCCTGCGCTACTTCAACAGC |
| DGC0995 prom-Rev | CCGGAATTCCACGATGCCATTTCCGCC |
| DGC1008 prom-For | CGCGGATCCCGTCAACGCATGATGATT |
| DGC1008 prom-Rev | CCGGAATTCTGTTATTCGCCTATTGCT |
| DGC1023 prom-For | CGCGGATCCTGGCGTTTAGCGATAACG |
| DGC1023 prom-Rev | CCGGAATTCAGTGATTTCCCAAGTAAA |
| DGC1024 prom-For | CGCGGATCCTTTCGCTTAACGACTGAC |
| DGC1024 prom-Rev | CCGGAATTCGTCCGCTCCTAAATTCCA |
| DGC1089 prom-For | CGCGGATCCATCCTTTGTCTCTGGTGT |
| DGC1089 prom-Rev | CCGGAATTCACGTCTGAACCCTGTAAC |
| DGC1854 prom-For | CGCGGATCCGAAAAGCCCTATACCGCG |
| DGC1854 prom-Rev | CCGGAATTCGCAAGAATCCAGCTGCGC |
| DGC2196 prom-For | CGCGGATCCTGAGGCGTTCCACAGTGA |
| DGC2196 prom-Rev | CCGGAATTCGTGCATCCCTGCTTCGAA |
| DGC2242 prom-For | CGCGGATCCCGGATGAATTTGCTTTAG |
| DGC2242 prom-Rev | CCGGAATTCTTTAAGGTGAGCCTGACA |
| DGC2334 prom-For | CGCGGATCCAGGTGTTGGCGCGCAAGC |
| DGC2334 prom-Rev | CCGGAATTCAACTTCTCCAGGCCACAT |
| DGC2465 prom-For | CGCGGATCCGCGCATAGTAGCAACGCC |
| DGC2465 prom-Rev | CCGGAATTCGCGCAATGCTCGCGAAAT |
| DGC2697 prom-For | CGCGGATCCGCTTTCCCAGCCAGGCCG |
| DGC2697 prom-Rev | CCGGAATTCGGAAACTTCCTCCGGGGG |
| DGC2884 prom-For | CGCGGATCCGTTAAATCACTTCAAGGG |
| DGC2884 prom-Rev | CCGGAATTCCTTATTTGCTTCCATTGC |
| DGC3006 prom-For | CGCGGATCCGGCTGGCACTTAAGTAAG |
| DGC3006 prom-Rev | CCGGAATTCGGTCGGCTGATGGAGAGG |
| DGC3134 prom-For | CGCGGATCCATCCAAAATGAAACTTTA |
| DGC3134 prom-Rev | CCGGAATTCGTGAAAACCTCAAAAGAG |
| DGC3217 prom-For | CGCGGATCCCTGTCCTAAACCTGACTC |
| DGC3217 prom-Rev | CCGGAATTCAGTGGTCGGAGCTCTTGA |

| | |
|---|---|
| DGC3247 prom-For | CGCGGATCCACAATACTTCTCATCTTG |
| DGC3247 prom-Rev | CCGGAATTCTATTATTCTCGTGACAGC |
| DGC3482 prom-For | CGCGGATCCCCGGTGCCGATCTCATTT |
| DGC3482 prom-Rev | CCGGAATTCGCGGTTACTCTTATTAAT |
| DGC3621 prom-For | CGCGGATCCGGCCATTTTACGACGCCA |
| DGC3621 prom-Rev | CCGGAATTCAACGCGCCGGCCTTAGTG |
| DGC4070 prom-For | CGCGGATCCGCAATTCCGCTTGCAGGG |
| DGC4070 prom-Rev | CCGGAATTCGCACGGGAAGTATCAGGA |
| DGC2884_AADEF For | CTG GTC GCC CGA TTA **GCC GCC** GAT GAG TTT GCC ATG |
| DGC 2884_AADEF Rev | CAT GGC AAA CTC ATC **GGC GGC** TAA TCG GGC GAC CAG |
| DGC2884 For (BamHI) | CGCGGATCCATGAAATTAGAAAATTCAATCAAC |
| DGC2884_noTM For (XbaI) | GC TCT AGA ATG AGT GAG TTT CTC CAT CGC |
| DGC2884 Rev (HindIII) | CCCAAGCTTTCATATCCGCACCTTACTCATTCC |
| GWdgc2884 For | CACCGTTAAATCACTTCAAGGG |
| GW DGC2884_noTM For | CACC ATG AGT GAG TTT CTC CAT CGC |
| GWdgc2884 Rev | TATCCGCACCTTACTCATTCC |

Underlined portions of primer sequences correspond to restriction enzyme sites: BamHI (For) and EcoRI or HindIII (Rev). Bold letters represent sites targeted for site-directed mutagenesis.

```
CLUSTAL O(1.2.4) multiple sequence alignment


Pantoea_DGC2884      -MKLENSINDKSSIRNKLRKISTINSAVILLLCWLLLSSTSLLFIKSYEKRNLELIAATL          59
PA01_TpbB            MNRRRRYTGSNPSLRRVLYRAHLGVALVAVFTAGLAVTLVGLLTLRAYADPNQQLIARSI          60
MG1655_DgcN          MMDNDNSLNKRPTFKRALRNISMTSIFITMMLIWLLLSVTSVLTLKQYAQKNLALTAATM          60
                          .  ... :::. * .       : ::   * :: ..:* :: * . *  * * ::

Pantoea_DGC2884      STTLTAATVFEDSYDAHNKIARLGNEGMFDSAKLVTDDQTVLVDWHDQQEQTG-WYPRLL          118
PA01_TpbB            SYTVEAAVVFGDAQAAEESLALIASSEEVSSAIVYDRQGQPLASWHRESTGPLHLLEQQL          120
MG1655_DgcN          TYSLEAAVVFADGPAATETLAALGQQGQFSTAEVRDKQQNILASWHYTRKDPGDTFSNFI          120
                     : :: **.** *.  * :.:* :...  ..:* :   :   *..**           . :

Pantoea_DGC2884      REWIYLKPFSVNIIHSETNVGTLTLEGTVTGAADFIRYSLMILTSGMLCVLIISFLLSEF          178
PA01_TpbB            AHWLLSAPTEQPILHDGQKIGSVEVKGSGGSLLRFLLTGFAGMVLCLLLTALGAFYLSRR          180
MG1655_DgcN          SHWLFPAPIIQPIRHNGETIGEVRLTARDSSISHFIWFSLAVLTGCILLASGIAITLTRH          180
                      .*:   *    * *.  .:* : : .   .   *:  .:  :.  :* .   :: *:.

Pantoea_DGC2884      LHRGILASLRNITSSIHYVIHSGDFSLRIPESPTREFQLFSNDLNSLLTEMQTLQASLMR          238
PA01_TpbB            LVRGIVGPLDQLAKVAHTVRRERDFEKRVPEAGIAELSQLGEDFNALLDELESWQARLQD          240
MG1655_DgcN          LHNGLVEALKNITDVVHDVRSNRNFSRRVSEERIAEFHRFALDFNSLLDEMEEWQLRLQA          240
                     * .*::  * :::.  * *  . :*. *: *    *:  :. *:*:** *::  *  *

Pantoea_DGC2884      DNRTLAAKALEDPLTGLANRAAFVARLTQLL-DQQLAKERFVLLFLDGDRFKSINDNWGH          297
PA01_TpbB            ENASLAHQAHHDSLTSLPNRAFFEGRLSRALRDASEHREQLAVLFIDSDRFKEINDRLGH          300
MG1655_DgcN          KNAQLLRTALHDPLTGLANRAAFRSGINTLM-NNSDARKTSALLFLDGDNFKYINDTWGH          299
                     .*  *   * .* **.* *** * . :.  : : .  ::  .:**:*.*.** ***  **

Pantoea_DGC2884      AAGDEVLKAIGSRLSTLAYKQDLVARLGGDEFAMLISSRASDAQLQLLMQDISESISQKI          357
PA01_TpbB            AAGDTVLVNIAMRIRGQLRESDLVARLGGDEFAVLLAPLASGADALRIADNIIASMQAPI          360
MG1655_DgcN          ATGDRVLIEIAKRLAEFGGLRHKAYRLGGDEFAMVLYDVQSESEVQQICSALTQIFNLPF          359
                     *:** **  *. *:       . . ********:::    * ::   : . :   :.  :

Pantoea_DGC2884      IIAEGTPISTSVTIGYAWS-QHGDTVESILERADMNMYKNKGMSKVRI------------          404
PA01_TpbB            RLSDGSTVSTSLTIGIALYPEHADTPAALLHDADMAMYIAKRQARGSRRLAELNDPRILQ          420
MG1655_DgcN          DLHNGHQTTMTLSIGYAMTIEHASAEK-LQELADHNMYQAKHQRAEKL--VR--------          408
                      : :*   : :::** *   :*..:   : . **  **   *

Pantoea_DGC2884      ---------------          404
PA01_TpbB            EEKEIDSATPEAPPK          435
MG1655_DgcN          ---------------          408
```

A

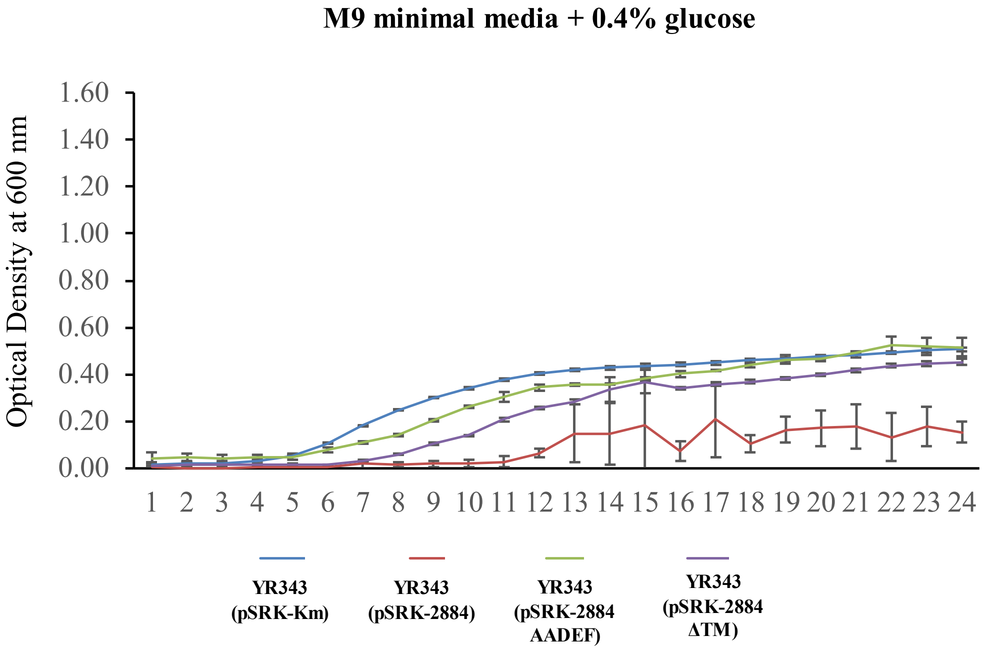
M9 minimal media + 0.4% glucose
Optical Density at 600 nm
1.60
1.40
1.20
1.00
0.80
0.60
0.40
0.20
0.00
1 2 3 4 5 6 7 8 9 10 11 12 13 14 15 16 17 18 19 20 21 22 23 24
YR343 (pSRK-Km)
YR343 (pSRK-2884)
YR343 (pSRK-2884 AADEF)
YR343 (pSRK-2884 ΔTM)

B

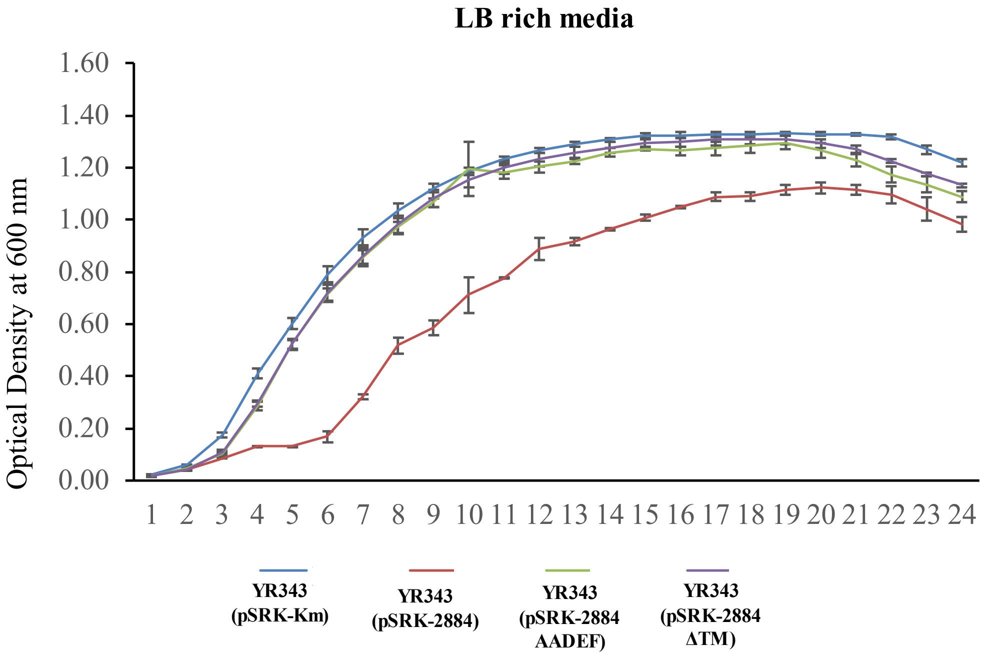
LB rich media
Optical Density at 600 nm
1.60
1.40
1.20
1.00
0.80
0.60
0.40
0.20
0.00
1 2 3 4 5 6 7 8 9 10 11 12 13 14 15 16 17 18 19 20 21 22 23 24
YR343 (pSRK-Km)
YR343 (pSRK-2884)
YR343 (pSRK-2884 AADEF)
YR343 (pSRK-2884 ΔTM)

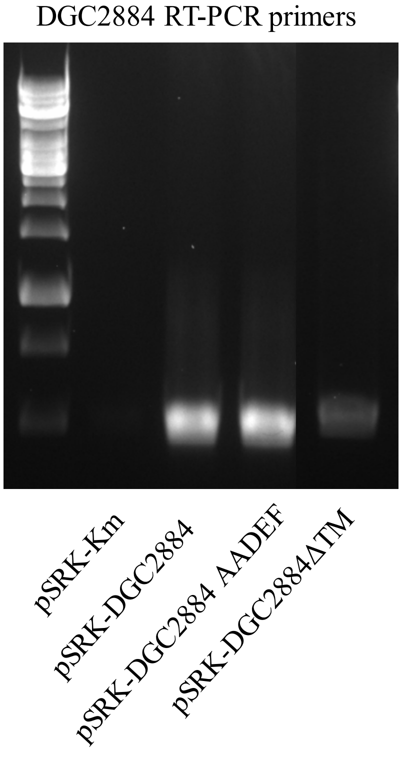
DGC2884 RT-PCR primers
pSRK-Km
pSRK-DGC2884
pSRK-DGC2884 AADEF
pSRK-DGC2884ΔTM

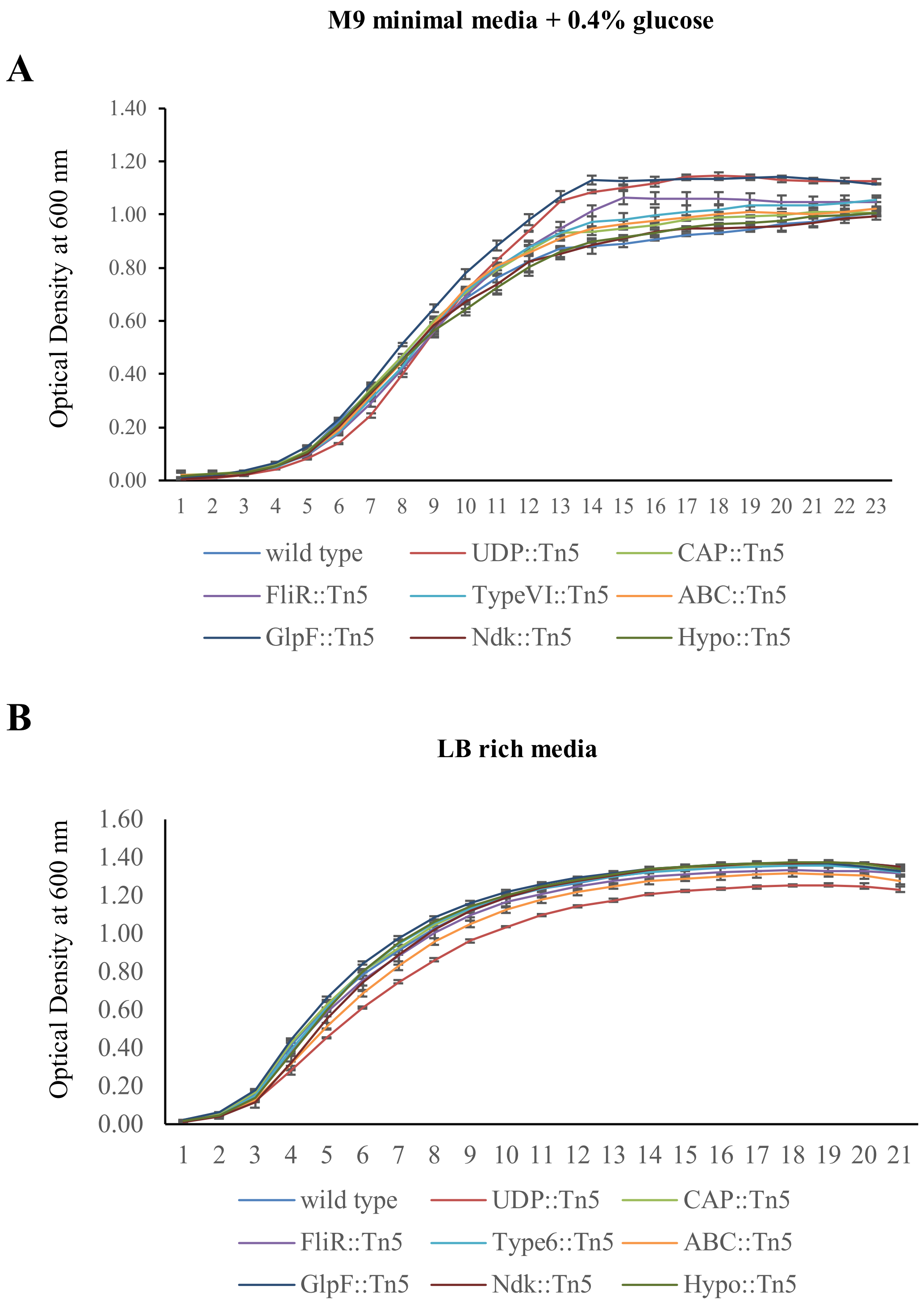

M9 minimal media + 0.4% glucose
A
Optical Density at 600 nm
1.40
1.20
1.00
0.80
0.60
0.40
0.20
0.00
1 2 3 4 5 6 7 8 9 10 11 12 13 14 15 16 17 18 19 20 21 22 23
wild type
UDP::Tn5
CAP::Tn5
FliR::Tn5
TypeVI::Tn5
ABC::Tn5
GlpF::Tn5
Ndk::Tn5
Hypo::Tn5
B
LB rich media
Optical Density at 600 nm
1.60
1.40
1.20
1.00
0.80
0.60
0.40
0.20
0.00
1 2 3 4 5 6 7 8 9 10 11 12 13 14 15 16 17 18 19 20 21
wild type
UDP::Tn5
CAP::Tn5
FliR::Tn5
Type6::Tn5
ABC::Tn5
GlpF::Tn5
Ndk::Tn5
Hypo::Tn5

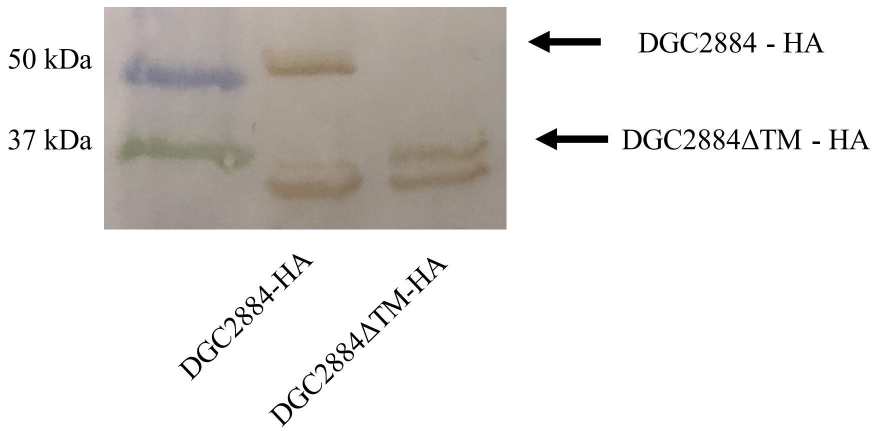
50 kDa
37 kDa
DGC2884 - HA
DGC2884ΔTM - HA
DGC2884-HA
DGC2884ΔTM-HA

Original Gel Image

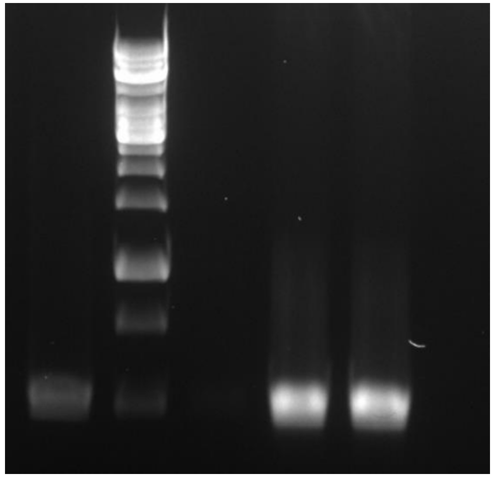


Original unedited western blot (3 sets)

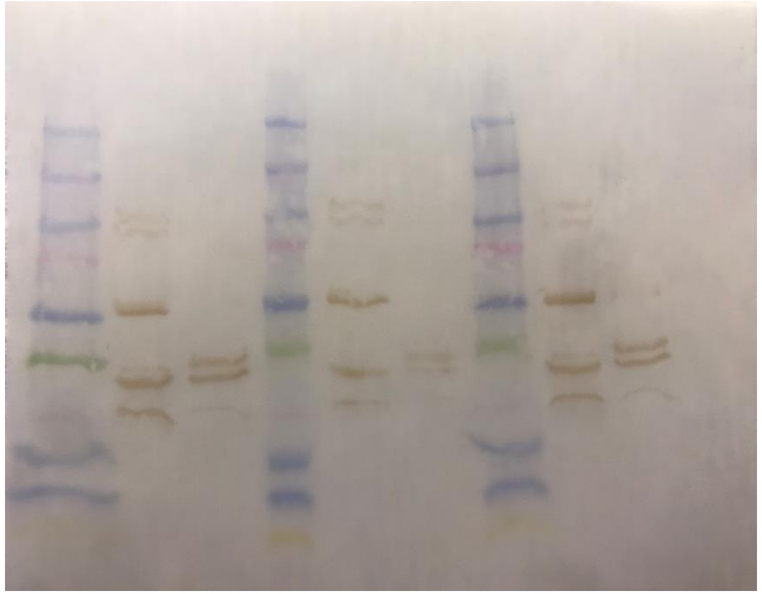
